# Cancer cells transfer invasive properties through microRNAs contained in collagen-tracks

**DOI:** 10.1101/2025.01.25.634857

**Authors:** Léa Normand, Lucile Rouyer, Elodie Richard, Nathalie Allain, Julie Giraud, Sylvaine Di-Tommaso, Cyril Dourthe, Anne-Aurélie Raymond, Jean-William Dupuy, Kevin Moreau, Richard Iggo, Gaetan McGrogan, Nathalie Dugot-Senant, Anthony Bouter, Sisareuth Tan, Alexandre Favereaux, Violaine Moreau, Reini F. Luco, Manon Ros, Frédéric Saltel

## Abstract

Invasion is a prerequisite for metastasis formation. During tumor development, the extracellular matrix (ECM) is remodeled in part through overexpression of type I collagen, increasing tumor microenvironment stiffness, and facilitating cancer dissemination. During breast cancer cell migration, we observed membrane debris left behind, attached to the collagen fibrils, along the migration path. We named these structures collagen-tracks. These collagen-tracks can be deposited in 3D matrices *in vitro* and *in vivo* and their formation is stimulated by the interaction between the ECM and matrix receptors, such as the discoidin-domain receptor (DDR1). However, they are different from structures already known to be involved in cell-cell communication such as exosomes and migrasomes, due to their specific nucleic acid and protein contents. When deposited by highly invasive breast cancer cells, internalized collagen-tracks reprogram non-invasive cells into highly pro-metastatic ones by inducing a partial epithelial–mesenchymal transition (EMT). This cell reprogramming is dependent on specific miRNAs present in the collagen-tracks, that are necessary to promote ECM degradation, increase cell motility and invasiveness. Collagen-tracks thus represent a new form of cell-cell communication important for driving tumor invasion that could be targeted to prevent metastasis.

## Introduction

Breast cancer is the most commonly diagnosed cancer in women, with the highest cancer-related mortality^1^. This is partly due to its heterogeneity, with multiple subgroups requiring specific treatments, but above all to metastasis formation. Tumor microenvironment (TME) remodeling includes extracellular matrix (ECM) accumulation associated with an increase in tissue stiffness. This element is a crucial driver of metastasis that is associated with poor prognosis^2,3^. In particular, ECM contains a dynamic collagen-rich meshwork, in which type I collagen is the most abundant matrix protein. Type I collagen is known to have multiple functions during tumor progression, notably promoting cell invasion and metastasis^4,5^. To achieve this, collagen interacts with cell surface receptors that sense physical signals from the ECM and activate intracellular signaling pathways^6–8^. Furthermore, fibrillar type I collagen is overexpressed in many cancers, including breast cancer, leading to its accumulation, increased cross-linking and alignment, promoting ECM stiffness, cell invasion and poor prognosis^2,9–12^ ^2,5,13^.

During migration and throughout cancer progression, cells strongly adhere to type I collagen, notably through adhesive structures like focal adhesions and invadosomes, that contain collagen receptors such as β1 integrins and discoidin domain receptors (DDRs)^14^. Both types of receptor are overexpressed in breast cancer, where they promote cell migration, ECM degradation and metastasis formation^4,15–17^.

During cell migration through dense ECM, cancer cells are subjected to physical stresses leading to cell membrane disruptions. Adhesion sites can be torn from the plasma membrane, leaving a trail of membrane debris on the ECM along the migration path. Few studies have analyzed these debris, which are left behind as cells retract their trailing edge^18,19^. Invasive melanoma and breast cancer cells have been shown to release integrins and CD44-enriched fragments during cell invasion in collagen matrices^20,21^. Such fragments called collagen-tracks may facilitate the reorientation and migration of the other cells along the preexisting path^22^. However, the molecular content of collagen-tracks and their biological role in cell invasiveness and tumor progression remain to be determined.

Here, we prove that, similar to the extracellular vesicles (EVs) released by cancer cells, these fragments also play a role in cell-cell communication and tumor progression^23,24^. EVs encompass exosomes, microvesicles, migrasomes and oncosomes that differ in size, molecular composition and mechanism of biogenesis^25^. EVs can be internalized and modify the gene expression profile, phenotype and properties of surrounding cells in multiple types of cancers. They contain proteins, RNAs and microRNAs that, once taken up by neighboring cells, can have functional effects by activating pro-tumoral signaling pathways, leading to increased tumor growth, cancer cell migration, ECM degradation and invasion^26^. Here, we demonstrate that collagen-tracks are different from well-known EVs in their protein, RNA and microRNA compositions. Their formation correlates with strong ECM-matrix receptor interactions mediated by receptors like DDR1 and integrins. Finally, we show that collagen-tracks released by migrating tumor cells are taken up by surrounding cells, modifying their gene expression profile and behavior. Unlike exosomes, their effect is strictly local because they are stably attached to fibrillar collagen, a fixed element in the ECM. Furthermore, collagen-tracks formed by aggressive breast cancer cells induce invasive features in non-invasive cancer cells by promoting epithelial-to-mesenchymal transition (EMT), ECM degradation and invasion. This reprogramming is mediated by specific miRNAs present in the collagen-tracks.

Taken together, our results show that collagen-tracks formed during cell migration in a collagen fiber-rich TME play a role in cell-cell communication by promoting cancer cell invasiveness.

## Results

### Membrane vesicles containing DDR1 are deposited on collagen fibers during 2D and 3D breast cancer cell migration

During breast cancer cell migration in a type I collagen context, we noticed plasma membrane staining (Cell Mask) along collagen fibrils in the extracellular space, following the migration path (Fig. 1a). This could be explained by shed membrane forming deposits during cell migration ^4,27,28^. As we observed the same pattern when staining for the membrane receptor DDR1, we named these DDR1-positive membrane structures in the cell migration path "collagen-tracks" (Fig. 1b, Fig. S1a)^29,30^.

**Fig. 1:**
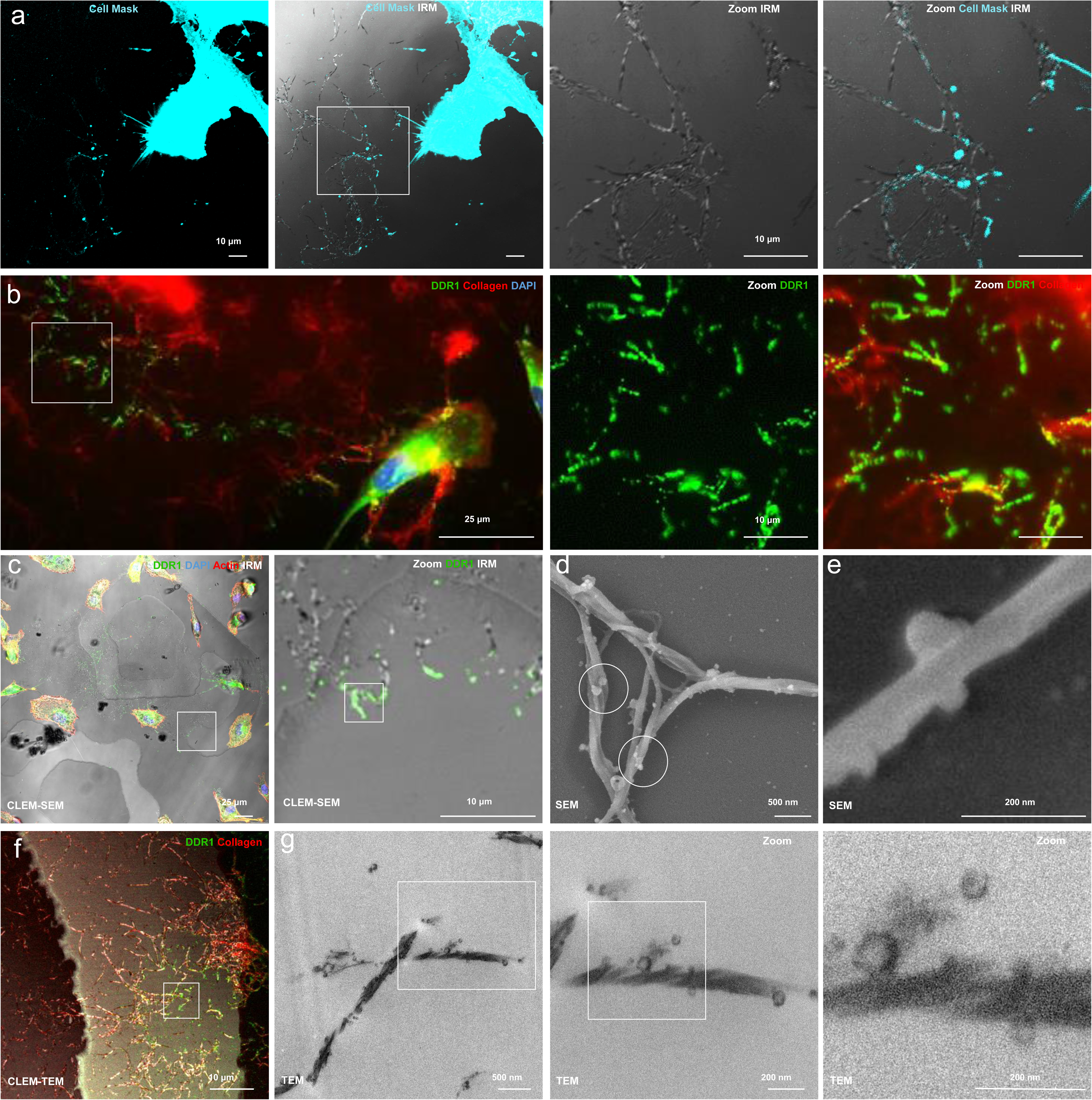
Membrane vesicles containing DDR1 are deposited on collagen fibers during breast cancer cell migration in 2D and 3D. **(a**) Immunofluorescence images of MDA-DDR1-GFP cells seeded on type I collagen and imaged for cell membrane (Cell Mask, in cyan, collagen imaged by IRM in grey). Scale bars, 10 μm. **(b)** Immunofluorescence images of MDA-DDR1-GFP cells seeded on type I collagen (in red) and stained for DDR1 (in green). Scale bars, 25 µm and 10 µm. **(c)** CLEM-SEM image of collagen-tracks after MDA-DDR1-GFP cells migration along type I collagen. DDR1 is stained in green, actin in red, nuclei in blue and collagen is imaged by IRM. Scale bars, 25 μm and 10 μm. **(d)** SEM image of collagen-tracks stuick on collagen fibers (white circles). Scale bar, 500 nm. **(e)** SEM image of zoomed collagen-tracks on one collagen fiber. Scale bar, 200 nm. **(f)** CLEM-TEM images of collagen-tracks after MDA-DDR1-GFP cells migration along type I collagen. DDR1 is stained in green, collagen in red. The square highlights the regio analysed in TEM in (g). Scale bar, 10 μm. **(g)** TEM images of collagen-tracks on type I collagen. Scale bars, 500 nm and zoom: 200 nm.

We analyzed the ultrastructure of collagen-tracks with multiple microscopy approaches. First, we performed correlative light and scanning electron microscopy (CLEM-SEM) on type I collagen incubated with MDA-MB-231 cells stably expressing DDR1 fused to GFP (MDA-DDR1-GFP cells) or, as a control, the same ECM incubated with conditioned media from MDA-DDR1-GFP cells (Fig. 1c-e and Fig. S1b). Using both SEM and transmission electron microscopy (TEM), we found that collagen fibers decorated with vesicle-like structures were only observed in regions positive for DDR1-GFP staining and were absent in control conditions (Fig. 1d-g and Fig. S1b, c). The size and shape of the vesicle-like structures was heterogeneous, with an average size of 50 ± 30 nm (Fig. S1d). To analyze their formation, we performed video time-lapse imaging using MDA-DDR1-GFP cells seeded on type I collagen fibers. We could show collagen-track formation on collagen fibers during cell detachment and migration (Movies S1, S2 and Fig. S1e-g). These live cell experiments demonstrated the large amount of material deposited by cells during migration, and confirmed the fact that these structures are torn from the membrane during migration. Collagen-tracks persist for at least 12 days after removing the cells from the ECM (Fig. S1h), and their number increases with time (Fig. S1i). To investigate whether migration is required for collagen-track formation, we blocked migration with a myosin II inhibitor or an actin polymerization inhibitor (cytochalasin D). Both drugs strongly reduced collagen-track formation, showing the importance of cell migration in their formation (Fig. S1j).

We then wanted to verify whether collagen-tracks can be found in 3D cultures and in tumors *in vivo*. First, we produced a dense fibrillar collagen gel in Boyden chambers and seeded MDA-DDR1-GFP cells on top of the gel. After induction of cell invasion by a serum gradient^4^, confocal z-stacks revealed the presence of cell-free collagen fibers decorated with DDR1 inside the gel (Fig. 2a), showing that collagen-tracks are also formed after migration in 3D culture. To test for their presence *in vivo*, we combined a xenograft model with a multiplex immunofluorescence approach. After intradermal implantation of MDA-DDR1-GFP cells expressing Caax-mCherry to visualize the cell membrane, we waited for 15 days after cell injection for tumor development and local invasion (Fig. 2b). Tumors were then removed together with the surrounding healthy tissue to collect cells that had started to invade (Fig. 2c). Labeled tumor cells were observed at a distance from the tumor, corresponding to local invasion (Fig. 2d-f). We were able to identify collagen-tracks *in vivo* by visualizing fibrillar collagen using second harmonic imaging, coupled with membrane co-labelling with DDR1-GFP and Caax-mCherry at a distance from the cell body, confirming the existence of these structures *in vivo* (Fig. 2e, f).

**Fig. 2:**
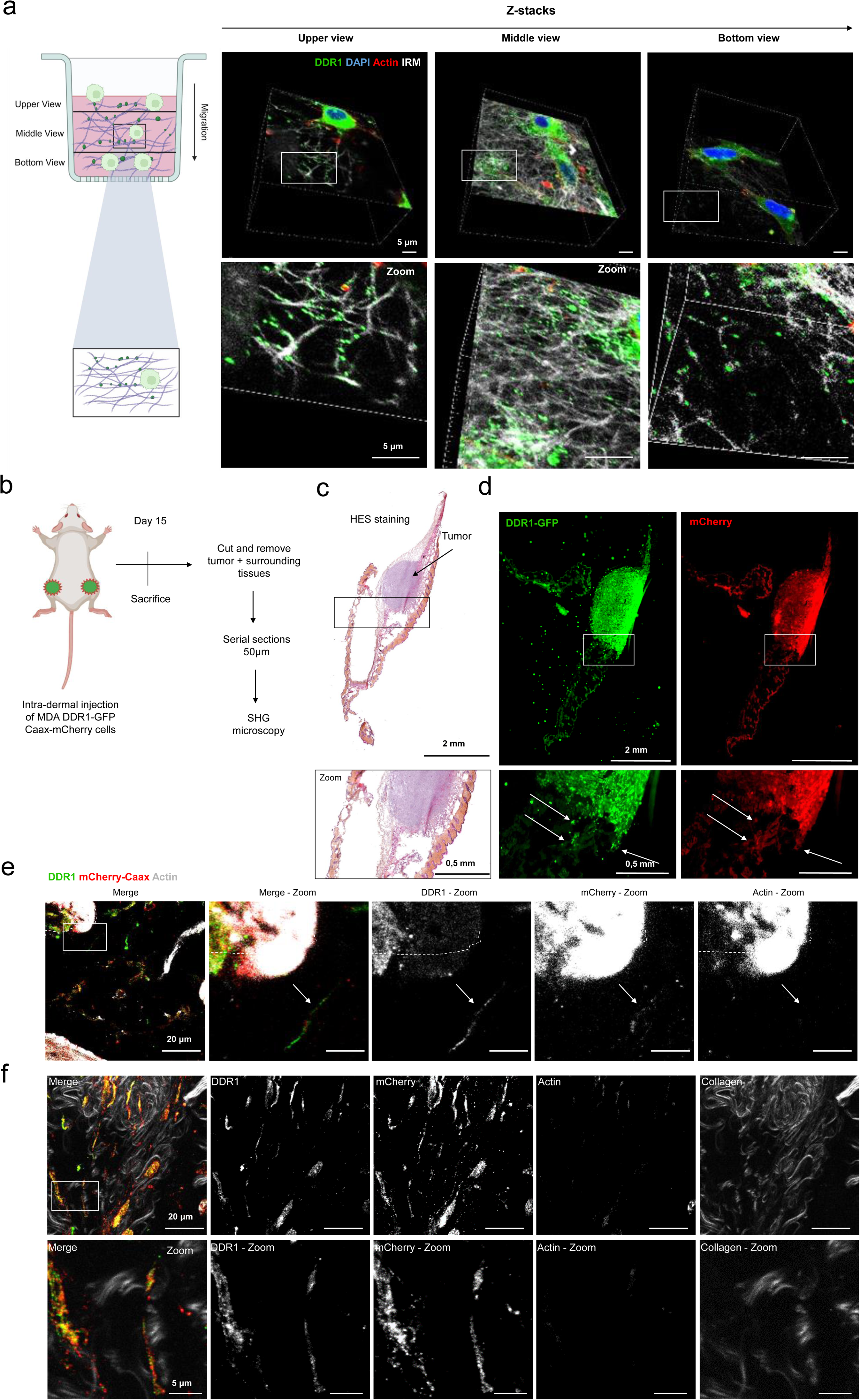
Tracks can be found in 3D and *in vivo*. **(a)** Experimental workflow for 3D collagen gel and representative 3D gel image (left panel). Immunofluorescence images of the 3D collagen gel (right panel), upper view (on the left) to lower view (on the right). Collagen-tracks are stained with DDR1 (green), actin (red), nucleus (DAPI) and collagen observed by IRM (grey). Scale bars, 5 µm. **(b)** Experimental workflow of intradermal injection of MDA-DDR1-GFP cells in mice and the following processing of tumors. **(c)** HES images of a mouse tumor. Scale bar: 2 mm, zoom: 0,5 mm. **(d)** Immunofluorescence images of the mouse tumor in (e). DDR1 is stained with GFP in green and cell membrane is stained with Caax-mCherry in red. Scale bar, 2 mm, zoom: 0,5 mm. **(e)** Immunofluorescence images of collagen-tracks after *in vivo* experiment described in (d). DDR1 is stained in green, cell membrane in red and actin in grey Scale bar, 20 µm. The dash line surrounds the cell and the arrow show the presence of collagen-tracks outside the tumor. **(f)** Immunofluorescence images of collagen-tracks after *in vivo* experiment described in (d). DDR1 is stained in green, the cell membrane in red, actin in grey and collagen is imaged by second harmonic generation microscopy. Scale bar, 20 µm, zoom: 5 µm.

Taken together, these results demonstrate that collagen-tracks are deposited on type I collagen fibers during cell migration and can be formed in 3D culture and *in vivo*.

### Collagen-track formation is stimulated by the interaction between collagen fibers and collagen receptors

To test whether collagen-track formation is specific to fibrillar collagen I, we seeded MDA-DDR1-GFP cells on other ECM components and analyzed collagen-track formation. MDA-DDR1-GFP cells seeded on non-fibrillar type I collagen, gelatin, matrigel or fibronectin, which stimulates migrasome formation^31^, did not form collagen-tracks (Fig. 3a, Fig. S2a). However, we did observe them when the cells were seeded on type III collagen fibers, another DDR1 ligand, showing that these structures are specifically formed on fibrillar collagens (Fig. 3a). Since collagen crosslinking is known to improve cell adhesion and dissemination in breast cancer^9,12^, we tested whether it affects collagen-track formation. LOX-L2 treatment, which increases collagen crosslinking, enhanced collagen-track formation by 4-fold (Fig. 3b and c). Furthermore, collagen-tracks were observed in all of the mammary cell lines we analyzed, including MCF-7, MDA-MB-486 and BT549, primary breast cancer cells and even normal mammary MCF10A cells (Fig. 3d, Fig. S2b). Since there is a positive correlation between DDR1 expression and collagen-track formation, we tested whether the absence of DDR1 could inhibit collagen-track formation. DDR1 knockout did not totally block their formation, as shown by membrane staining with Cell Mask, indicating that DDR1 promotes but is not essential for their formation (Fig. S2c, d). Since β1 integrins are also known to interact with collagen fibers^14^, we tested the impact of integrin activation using manganese (Mn2^+^)^32^ on collagen-track promotion. This showed an additive effect to DDR1 overexpression (Fig. 3e, f and Fig. S2e, f).

**Fig. 3:**
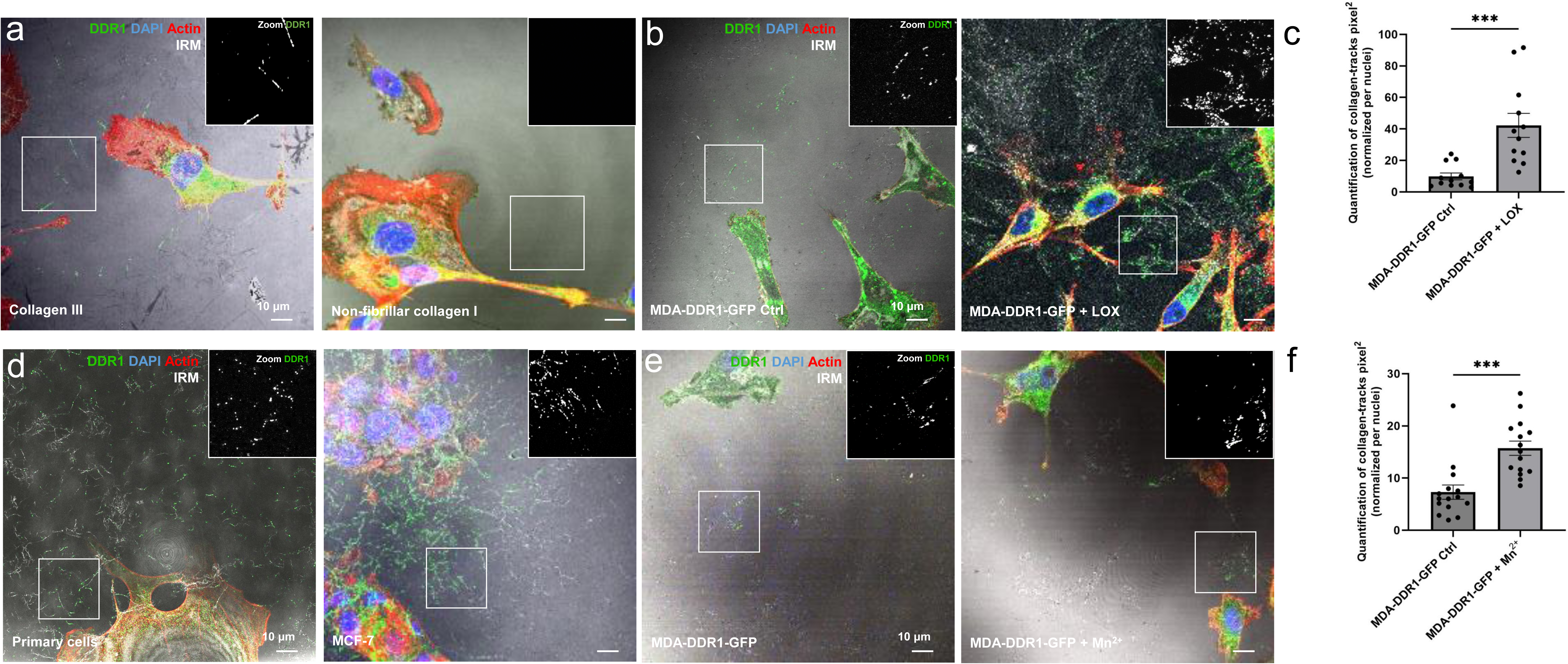
Collagen-track formation is stimulated by Collagen fibers-DDR1/integrin interaction. **(a)** Immunofluorescence images of MDA-DDR1-GFP cells seeded on collagen III and non-fibrillar collagen I matrices by MDA-DDR1-GFP cells. Cells were imaged by confocal microscopy. DDR1 is stained in green, actin in red, nuclei in blue and collagen is imaged by internal reflection microscopy (IRM) in grey. Scale bar, 10 μm. **(b)** Immunofluorescence images of collagen-tracks formation by MDA-DDR1-GFP cells on type I collagen and treated or not with LOX-L2. DDR1 is stained in green, actin in red, nuclei in blue and collagen imaged by IRM in grey. Scale bar, 10 µm. **(c)** Quantification of collagen-tracks formation by MDA-DDR1-GFP cells on type I collagen and treated or not with LOX-L2. Values represent the mean +/- SEM of n=3 experiment (with 5 images per condition). **(d)** Immunofluorescence images of basal primary breast cancer cells or MCF-7 cells seeded on type I collagen during 24h and stained for DDR1 in green, actin in red, nuclei in blue and collagen is imaged by IRM in grey. Scale bar, 10 μm. **(e)** Immunofluorescence images of collagen-track formation by MDA-DDR1-GFP cells on type I collagen and treated or not with Mn2+. DDR1 is stained in green, actin in red, nucleus in blue and collagen imaged by IRM in grey. Scale bar, 10 µm. **(f)** Quantification of collagen-track formation by MDA-DDR1-GFP cells on type I collagen and treated or not with Mn2+. Values represent the mean +/- SEM of n=3 experiments (5 images per experiment and per condition).

These data demonstrate that most mammary cells can form collagen tracks during migration on type I collagen. Taken together, these data highlight the importance and the complexity of the interaction between collagen fibers and their associated receptors in the formation of collagen tracks.

### Collagen-tracks have a novel protein composition

To characterize collagen-tracks, we first assessed their protein composition using a targeted approach based on well-known markers of EVs. We did not find exosome markers like CD63 and CD9, nor migrasome markers like TSPAN4, colocalizing with collagen-tracks (Fig. S3a), which aligns with the absence of collagen-tracks on fibronectin (Fig. S2a). However, some focal adhesion or invadosome markers were present on collagen-tracks, including CD44, Myosin IC, vinculin, β1 integrin and Tks5, but not cortactin or actin (Fig. S3b, c and Fig. 2e, f). These results demonstrate that collagen-tracks are distinct from known EVs and classical migrasomes.

To assess the full protein composition of this new type of EV, we performed mass spectrometry on decellularized collagen matrix decorated with collagen-tracks deposited by MDA-DDR1-GFP cells. As a control, we used collagen matrix incubated with conditioned medium from the same cells. MDA-DDR1-GFP cells were seeded overnight on type I collagen fibers to allow collagen-track formation, and then cells were removed by combined EDTA baths and laser microdissection (Fig. S4a, b)^33^. More than 200 proteins were specifically enriched in collagen-tracks, including DDR1, vinculin and β1 integrin, as expected (Fig. S3c and d, S4c, Table S1). Proteomic comparison with exosome and migrasome data sets, confirmed the uniqueness of collagen-tracks, with few shared proteins (Fig. S4d and Table S2). Gene set enrichment analysis (GSEA) confirmed that collagen-tracks are enriched in anchoring and adhesion molecules, validating the link with focal adhesion (Fig. S4e, f). We also identified proteins involved in membrane repair (AnxA1 and AnxA2), suggesting that membrane damage occurs during collagen-track deposition (Table S1). Finally, 54 RNA-binding proteins were seen, strongly suggesting that RNA is present in collagen-tracks (Fig. S4f).

### Collagen-tracks contain mRNAs and microRNAs

To dig deeper into the molecular composition of collagen-tracks, we performed an RNA sequencing analysis using the same conditions as for the proteomic analysis. Despite the low amount of material, we found that collagen-tracks contain mRNA, whereas we did not identify mRNA in the control condition using conditioned media. The volcano plot in Fig. 4a shows 32 mRNAs significantly identified in collagen-tracks (in red), many of which are involved in translation and protein degradation (Ingenuity Pathway Analysis, IPA) (Fig. 4c-d). Additionally, we found that among the 31 enriched mRNAs, 4 correspond to proteins that are also enriched in collagen-tracks, including RACK1, which is known to interact with transmembrane receptors like β1 integrin and promotes EMT^34^ (Fig. 4b).

**Fig. 4:**
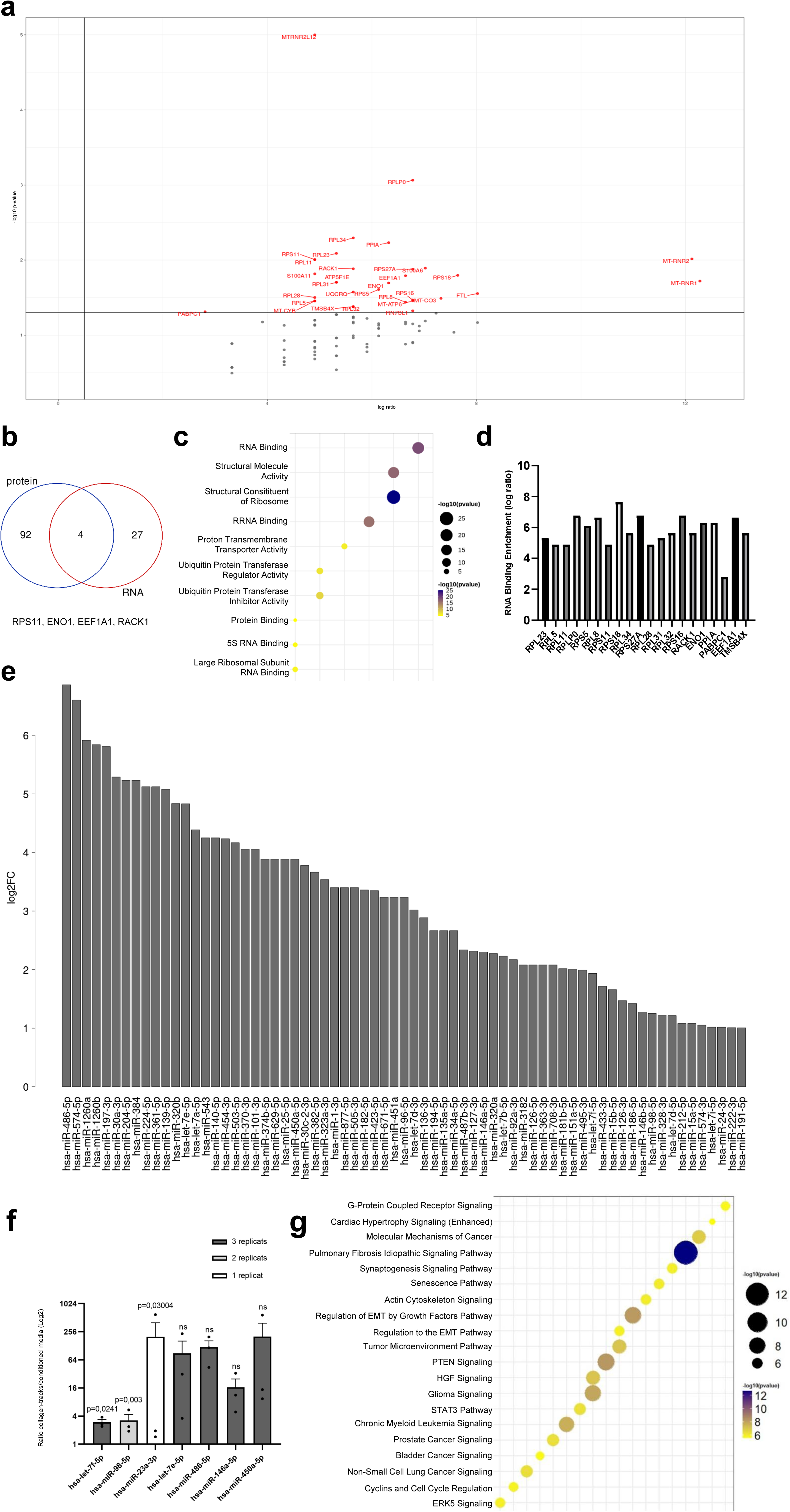
Collagen-tracks contain mRNAs and microRNAs. **(a)** Volcano map highlighting the 32 most expressed mRNAs enriched in collagen-tracks after RNA sequencing analysis **(b)** Venn diagram analysis of the 4 common proteins and mRNAs contained in collagen-tracks. **(c)** Bubble plot of the mRNAs related pathways. **(d)** Histogram of RNA binding RNA significantly present in collagen-tracks. Results are expressed in Log (ratio tracks/control). **(e)** Barplot of miRNAs enriched in collagen-tracks compared to control. **(f)** Histogram of the 7miRNAs significantly present in collagen-tracks and/or enriched into 1 to 3 replicates. Values represent the mean +/- SEM of n=3 independent experiments. **(g)** Bubble plot of the 7 miRNAs enriched and/or significantly present in collagen-tracks related pathways.

Since microRNAs (miRNA) are an important component of EVs, we performed miRNA sequencing and identified 70 miRNAs showing at least 2-fold in collagen-tracks compared to the control (Fig. 4e). Due to a high heterogeneity of miRNAs composition, we focused on seven miRNAs that were significantly upregulated in all three replicates (Fig. 4f). They include miRNA-23a, which is already known to be involved in the regulation of cell migration and invasion in breast cancer^35^. Using the miRDB database, we predicted the targets of these miRNAs, and performed an Ingenuity Pathway Analysis (IPA). As expected, pathways related to cancer, tumor microenvironment and actin signaling were identified (Fig. 4g). We also identified several pathways related to EMT and growth factor signaling, which goes in line with the known involvement of let-7f miRNA family in EMT^36,37^, let-7f-5p being the most significantly enriched miRNA in our analysis (Fig. 4e-g). These results demonstrate that collagen-tracks contain a substantial nucleic acid cargo notably involved in tumor microenvironment remodeling, EMT and cancer, suggesting that they might have the ability to modify the phenotype of cells that internalize them.

### Internalization of collagen-tracks induces a partial EMT and an invasive switch in recipient cells

To assess whether collagen-tracks can be internalized by other cells, we generated a cell membrane-labeled Caax-mCherry MCF-7 cell line. This less invasive breast cancer cell line was seeded on collagen decorated with MDA-DDR1-GFP collagen-tracks (Fig. S5a). Imaging of live and fixed MCF-7 cells revealed the presence of intracellular GFP, proving the internalization of DDR1-GFP collagen-tracks (Movie S3 and Fig. 5a, b). Quantification of the GFP signal showed that around 40% of MCF-7 cells internalize collagen-tracks after 72 hours (Fig. S5b). Transcriptomic analysis of MCF-7 cells grown for 72h on ECM decorated with MDA-DDR1-GFP collagen tracks identified 817 upregulated genes and 53 downregulated genes (Fig. 5c). Gene Ontology analysis of the upregulated genes highlighted terms associated with chemotaxis, ECM organization, cell adhesion, wound healing and migration, that together form a protein network related to ECM and cell migration (Fig. 5d, e). Since all of these terms are characteristic of EMT reprogramming, we compared the expression changes induced by collagen tracks with EMT signatures from the most recent EMT models annotated in the EMTome^38^. Almost 30% of all the changes in gene expression induced by collagen-tracks overlapped with changes characteristic of EMT (Fig. S5c and Table S3). These results were validated by RT-qPCR, western blot and immunofluorescence for well-known mesenchymal markers, such as vimentin and ZEB1, that were upregulated in MCF-7 cells after internalizing collagen tracks, whereas Snail was already expressed in control MCF-7 cells (Fig. S5d-f).

**Fig. 5:**
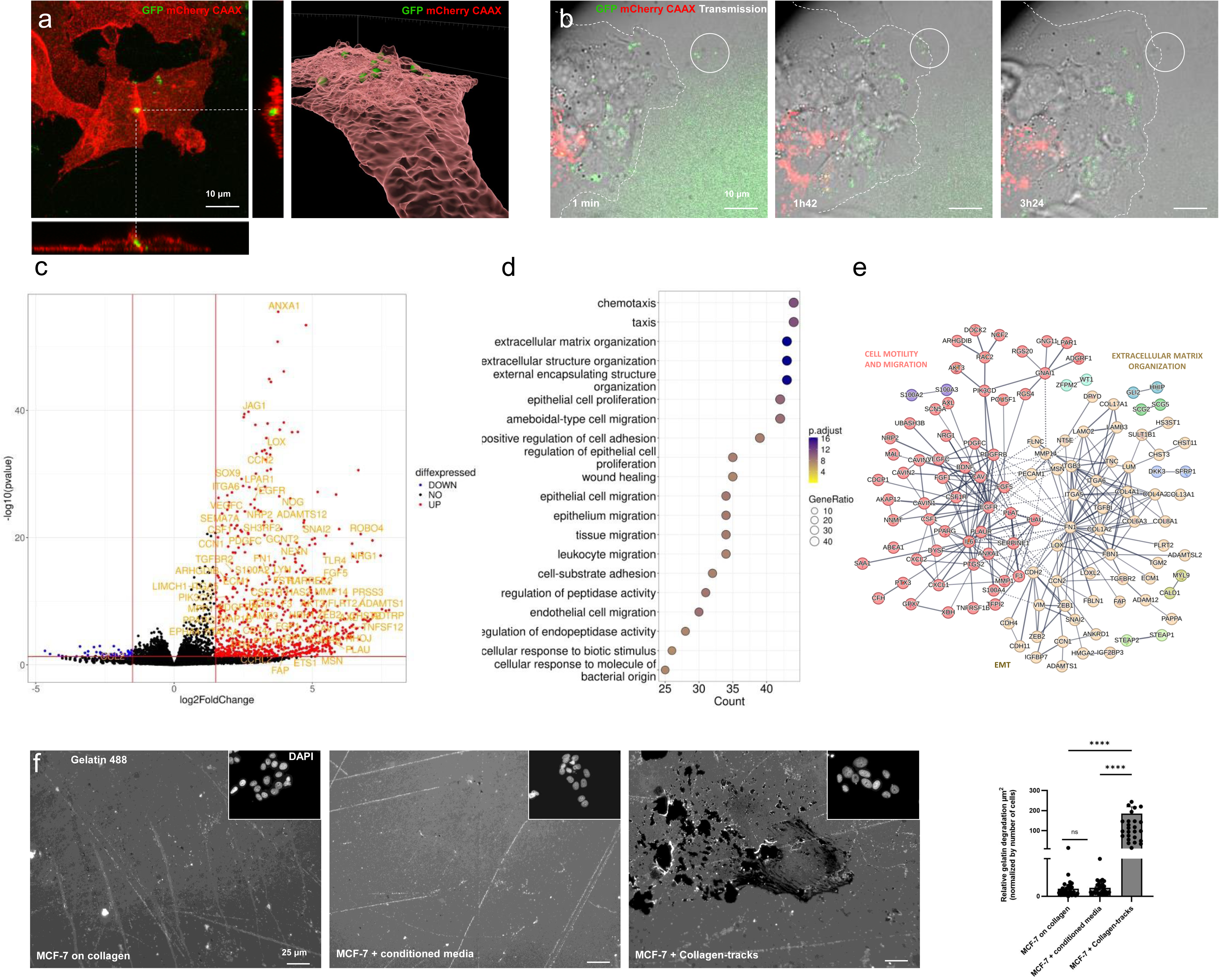
Internalization of collagen-tracks induces a partial-EMT and an invasive switch in recipient cells. **(a)** Immunofluorescence images with orthogonal views of MCF-7 cells that internalized collagen-tracks formed by MDA-DDR1-GFP cells. Cell membrane is stained with mCherry and collagen-tracks with GFP. Scale bar, 10 μm. On the right, the same image is represented in 3D using the Imaris software, the membrane is stained in red and collagen-tracks are represented by green spheres. **(b)** Images extracted from time lapse imaging showing the collagen-track internalization by MCF-7 cells. MCF-7 cells were seeded on a collagen matrix decorated with collagen-tracks. Cells were imaged by videomicroscopy for 15h. Collagen-tracks are observed with GFP stained in green and shown with white circle and cell membrane are observed with mCherry stained in red. Scale bar, 10 μm. **(c)** Volcano plot of transcriptomic analysis of MCF-7 internalizing collagen-tracks. Genes highlighted in yellow are upregulated and related to EMT. **(d)** Bubble plot of upregulated genes highlighted terms associated with ECM organization, cell adhesion, wound healing and migration of Gene Ontology enrichment analysis. **(e)** STRING analysis of upregulated genes associated into a protein network related to ECM and cell migration. **(f)** Left: Representative immunofluorescence images of ECM degradation properties of MCF-7 cells after collagen-tracks internalization compared to controls. Scale bar, 25 μm. Right: Quantification of ECM degradation properties of MCF-7 cells after being in contact with or without collagen-tracks. Values represent the mean +/- SEM of n=3 independent experiments (10 images per condition and per replicate) and were analyzed using One-way ANOVA test followed by bonferroni test p<0.0001.

Finally, since most of the changes in gene expression induced by collagen-tracks impact genes that are involved in cell migration (integrins, MMPs) and ECM remodeling (collagens, fibronectin) (Fig. 5e), we tested the impact of collagen-tracks on cell invasiveness by performing ECM degradation and cell migration assays in Boyden chambers (Fig. 5f and Fig. S5g, h). Using in-situ zymography, we showed that, after internalization of collagen-tracks, MCF-7 cells acquire the ability to degrade the ECM (Fig. 5f). This gain in invasiveness was dramatically reduced by inhibiting metalloproteinases (MMP) with the MMP inhibitor GM6001, which shows that this ECM degradation is MMP-dependent (Fig. S5g). Finally, Boyden chamber assays confirmed the gain in cell motility of MCF-7 cells after internalizing collagen-tracks, which further supports the capacity of collagen-tracks from highly invasive cells to reprogram recipient cells into highly invasive ones (Fig. S5h).

Taken together, these results demonstrate that the internalization of collagen-tracks laid down by invasive cells induces invasive characteristics and partial EMT in recipient cells.

### microRNAs contained in collagen-tracks promotes EMT of recipient cells

Since some of the miRNAs identified in collagen-tracks have been implicated in EMT regulation and induction of cell migration and invasion, we decided to test their role in MCF-7 reprogramming. Synthetic anti-microRNAs were used to block the effect of each of the seven miRNAs most consistently found in collagen-tracks laid down by MDA-DDR1-GFP cells (miR23a, miR-98, miR-146a, miR-450, miR-486, miRNA Let-7f and miRNA Let-7e). Briefly, the MDA-DDR1-GFP cells were transfected with individual anti-microRNAs for 5 days then seeded on collagen to lay down collagen-tracks. The MDA-DDR1-GFP collagen-track donor cells were then replaced with MCF-7 recipient cells on top of the collagen-track containing matrix. These MCF-7 were grown for 72h and we assessed the impact of miRNA inhibition in the donor cells on EMT reprogramming and cell invasiveness of recipient cells (Fig. 6a). miRNA inhibition did not affect MDA-DDR1-GFP cell migration or the formation of collagen-tracks (Fig. S6a). MCF-7 cells that internalized collagen-tracks in which only a single miRNA had been depleted using anti-miRNAs showed decreased expression of EMT markers such as vimentin, MMP14 and ZEB1, and less degradation of gelatin than control cells (Fig.6 b-d and Fig. S6b). This shows that these miRNAs are required for EMT reprogramming by collagen-tracks, elucidating the molecular mechanisms whereby collagen-tracks promote this invasive switch.

**Figure 6:**
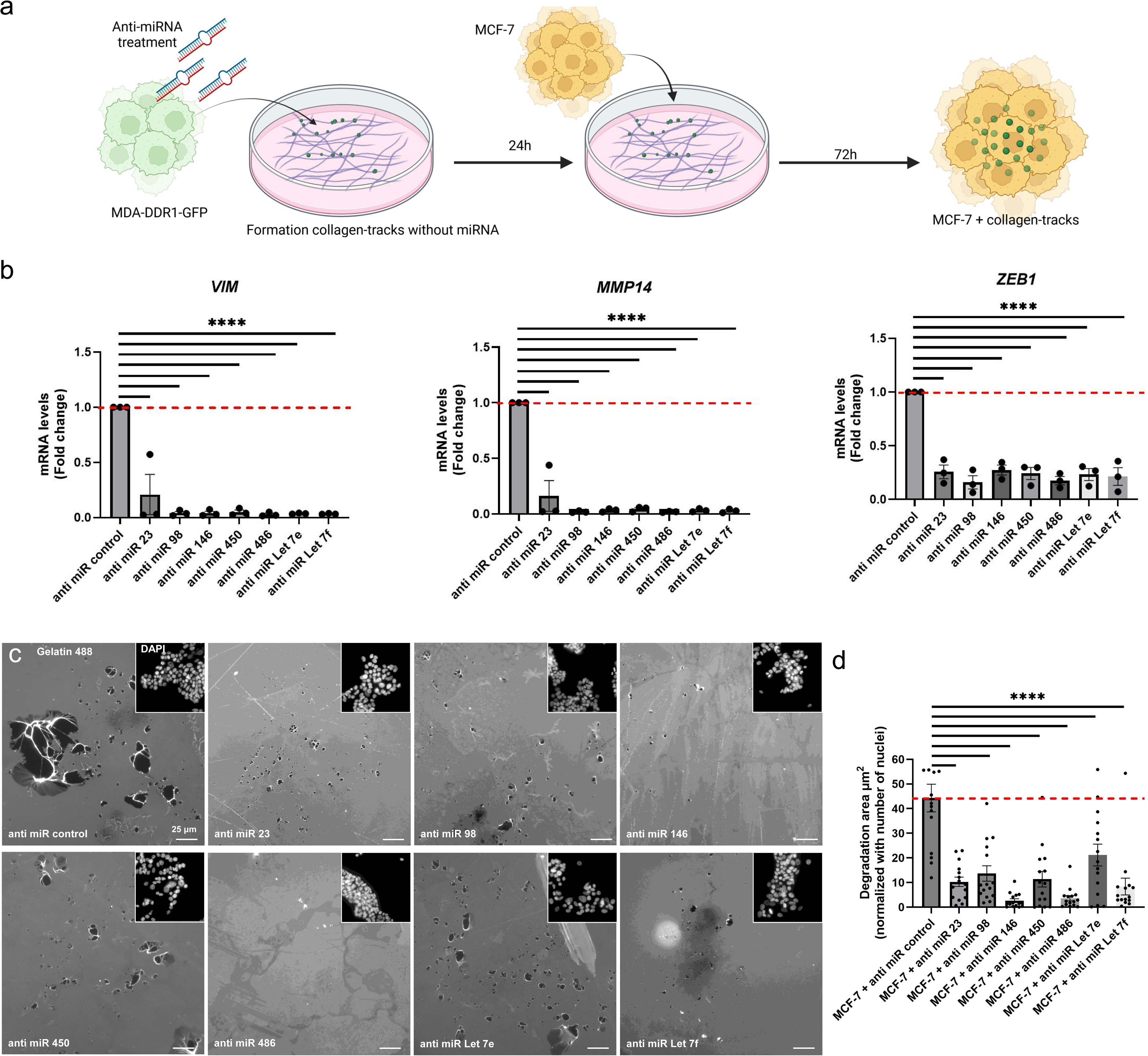
microRNAs contained in collagen-tracks promotes EMT of recipient cells. **(a)** Schematic procedure to form collagen-tracks without miRNAs of interest using miRNA inhibitors treatment. **(b)** mRNA expression levels of EMT genes in MCF-7 cells after internalization of collagen-tracks previously formed with anti-miRNAs treatment. The graph shows the quantification of *VIM, MMP14* and *ZEB1* mRNA expression levels normalized to 18S (Fold change). Values are expressed as the mean ± SEM of n=3 independent experiments. One-way ANOVA test followed by bonferroni test p<0.0001. **(c)** Representative immunofluorescence images of ECM degradation properties of MCF-7 cells after being treated with or without anti-miRNAs. Scale bar, 25 μm. **(d)** Quantification of ECM degradation properties of MCF-7 cells after being treated with or without anti-miRNAs. Values represent the mean +/- SEM of n=3 independent experiments (5 images per condition and per replicate) and were analyzed using One-way ANOVA test followed by bonferroni test p<0.0001.

## Discussion

In this study, we deciphered the composition and function of collagen-tracks, membrane deposits released along collagen fibers by cancer cells during migration. These structures are involved in cell-cell communication during cancer progression. They can be internalized by recipient cells and reprogram them into more invasive cells through the transfer of miRNAs.

If we compare collagen-tracks with other extracellular vesicles such as exosomes or migrasomes, their size is comparable to that of nanovesicles like exosomes^39^. Moreover, collagen-tracks can be formed by other cell types, such as fibroblasts, and help, such as Little Thumb, other cells to find their way^22^. Collagen-tracks from cancer cells have a unique molecular composition, in terms of proteins and nucleic acids, that sets them apart from classical migrasomes and exosomes. They are negative for the exosome markers CD9 and CD63, and negative for the migrasome marker TSPAN4. Instead, we showed that the collagen receptor DDR1 is a marker for collagen-tracks, colocalizing with membrane staining on type I and type III collagen fibrils.

We have shown that collagen-track formation is linked to the adhesive properties of cancer cells. Indeed, DDR1 overexpression, integrin activation, collagen accumulation and collagen crosslinking are elements that promote collagen-track formation during migration. However, the precise molecular mechanisms governing collagen-track formation remain to be elucidated. We have already identified additional collagen receptors, such as β1 integrin and CD44 that are present in collagen-tracks and could be involved, alone or in combination, in their formation. However, the mechanisms and molecular actors that control cell adhesion are multiple and potentially redundant, which complicates the analysis of the requirements for collagen-track formation. Indeed, the molecular composition of adhesive structures is very dynamic and complex, with multiple receptors cooperating within the same structure. This probably explains why inhibiting the expression of individual receptors did not block collagen-track formation. In contrast, blocking cell migration did limit collagen-track formation, highlighting the crucial link between track formation and migration. Furthermore, we have previously showed that migration of MDA-MB-231 cells along type I collagen fibers leads to membrane damage^27^. The resulting lesions are repaired by the concomitant action of annexins 2, 5 and 6 ^27,40^. Interestingly, in the mass spectrometry analysis reported here, we identified two annexins, ANXA1 and ANXA2, that could participate in this process. We postulate that collagen-tracks arise when rapidly migrating cells do not have time to properly disassemble adhesion complexes at sites of strong adhesion (Table S1). This type of structure was already observed in cancer cells expressing constitutively activated integrins, like the N305T β3 integrin mutant^41^. Without efficient membrane repair, cancer cells cannot survive, migrate or invade, highlighting a potential therapeutic target in cells that generate collagen-tracks^27,40^.

DDR1 is an important marker for collagen-tracks. In MDA-MB-231 cells and more generally in cancer cells, DDR1 can be present at cell-cell junctions, in linear invadosomes, in lamellipodia and now in collagen-tracks^42^. Furthermore, it was reported that ADAM10 controls the activity and half-life of DDR1 by cleaving off its ectodomain^29^. In this matrix context, the extracellular domain of DDR1 is found attached to collagen fibrils. Here we have demonstrated that collagen-tracks are associated with the full length DDR1 using proteomic data and a DDR1-GFP construct. Interestingly, DDR1 expression in triple negative breast cancer tumors enforces alignment of collagen fibrils, inhibiting immune infiltration^43^.

Some breast cancer cells and tissues express DDR2, the other member of the discoidin domain receptor family^44^. Like DDR1, DDR2 may be a player in collagen-track formation, depending on the cell type. Rather than targeting the receptors, an alternative therapeutic option to inhibit collagen-track formation might be to reduce or inhibit cancer cell adhesion to the ECM. Since the expression and chemical modification of matrix elements and receptors differs between tumors, we expect the molecular composition of collagen-tracks to differ in ways that will require more study across multiple tumor types before embarking on clinical trials.

After being released along collagen fibrils in the tumor microenvironment, collagen-tracks can be taken up by neighboring cells, where they induce partial EMT, migration and invasion thanks to their biologically active cargo. Partial EMT is a plastic state in which cells coexpress epithelial and mesenchymal markers. The impact of collagen-tracks may go beyond the communication between cancer cells, and collagen-tracks could have a wider impact on all the cell types that constitute the microenvironment.

We have shown that miRNA plays a crucial role in the reprogramming function of collagen-tracks. However, this function could be more complex and involve other elements. We expect the protein and RNA cargo of collagen tracks to also participate in the reprogramming function. This cargo notably includes RACK1, a protein associated with adhesion receptor signaling and EMT (Table S1). Another constituent is annexin A1, a positive regulator of MMP-9 expression that induces invasion through activation of the NF-κB signaling pathway, and is known to be overexpressed in MDA-MB-231 cells compared to MCF-7 cells^45^. Moreover, in a previous study, we showed that the translation machinery is required for invadosome formation and activity^33^. In the same way, collagen-tracks contain some elements of the translation machinery such as EEF1A1 that may modify the phenotype of recipient cells. Indeed, we suspect that it is the combined action of multiple elements in the collagen-track cargo that confers invasiveness on recipient cells.

Our unpublished data suggest that the content of collagen-tracks will depend on cell type and ECM composition, thus it will be necessary to analyze the composition of collagen-tracks from invasive and non-invasive cancer cells, but also from normal cells such as fibroblasts. Indeed, we expect the nucleic acid composition of collagen-tracks to differ from one cell type to another. Consequently, the impact on recipient cells could also differ, potentially even resulting in an anti-tumoral effect. In addition to tumor cells, other cells from the TME may be able to form collagen-tracks, such as cancer associated fibroblasts (CAFs). CAFs actively remodel the ECM, secrete multiple biologically active peptides, and have a high affinity for collagens^22,46^. Together, these elements confer them the potential to be major users of the collagen-track pathway, and point to the existence of a new layer of interdependence in the microenvironment where the ECM serves as a platform for cell-cell communication regulating cancer cell invasion.

Finally, we show the presence of DDR1 associated with membrane along collagen fibers after cancer cell invasion in a mouse model, suggesting the presence of collagen-tracks *in vivo*. Whether these collagen-tracks promote aggressiveness in the peri-tumoral microenvironment remains to be further explored. As recently demonstrated for exosomes^47^, cancer cell-released collagen-tracks could also promote leader-follower behavior in 3D migration.

In summary, we have identified and characterized an extracellular vesicle entity formed upon cell migration through shedding of plasma membrane fragments at sites of tight adhesion of cells to the ECM. Contrary to exosomes, they appear to act locally, with collagen-tracks playing a role in cell-cell communication and the transfer of invasive properties to surrounding cells. Cancer-related collagen-tracks are thus a new player acting locally to drive tumor invasion and metastasis. They constitute an additional mechanism through which ECM can control tumor dissemination, highlighting the complexity of the intercellular communication processes happening in the TME during tumor progression.

## Materials and Methods

### Antibodies and reagents

Anti-DDR1 (5583S), anti-CD9 (13174) and anti-clathrin light chain (4796S) were purchased from Cell Signaling. Anti-β1 integrin (ab52971 and ab30394), anti-calnexin (ab22595), anti-GFP (ab6673) and anti-mCherry (ab183628) antibodies were purchased from Abcam. Anti-CD63 (SAB4301607) and anti-vinculin antibodies (V9131) were purchased from Sigma Aldrich. Anti-myosin 1C antibody (A6936) was purchased from Abclonal. Anti-Tks5 (sc-376211) and anti-CD44 (sc-7297) were purchased from Santa Cruz. Anti-cortactin antibody (05-180) was purchased from Millipore. Secondary antibodies for immunofluorescence IRDye680 Conjugated Goat anti-mouse (926-32220, LI-COR), IRDye680 Conjugated Goat anti-rabbit, (926-32221, LI-COR), IRDye800 Conjugated Goat anti-mouse (926-32210, LI-COR), IRDye800 Conjugated Goat anti-rabbit (926-32211, LI-COR), Hoechst (H21486), phalloidin (FP-BZ-9630) and 5-carboxy-X-rhodamine succinimidyl ester were purchased from Interchim. Cell Mask reagent was purchased from Invitrogen. Type I collagen (rat tail, 354236) was purchased from Corning, type III collagen (5021) was purchased from CellSystems, fibronectin (F1056) was purchased from Sigma Aldrich and matrigel was purchased from Dutscher (#356231). Human TSPAN4-GFP plasmid was purchased from GenScript (NM_001025239.1). Species-specific fluorescent far-red coupled secondary antibody for western blot IRDye 680CW goat (polyclonal) anti-rabbit IgG (H+L) (926-68070) or IRDye 800CW goat (polyclonal) anti-mouse IgG (H+L) (926-32210) were purchased from LI-COR. GM6001 (CC10) and manganese (M7899-500G) were purchased from Sigma Aldrich.

### Cell culture

MDA-MB-231, MCF-7, BT549, MDA-MB-468 and MCF-10A cells were obtained from American Type Culture Collection. MDA-MB-231 and MCF-7 cells were maintained in DMEM high glucose (Gibco) supplemented with 10% fetal bovine serum (FBS, Sigma Aldrich), at 37°C in a 5% CO_2_ incubator. In this study, we used MDA-MB-231 cells transduced with a lentiviral plasmid expressing the discoidin domain receptor 1 (DDR1) fused to GFP and the cell line was named MDA-DDR1-GFP. BT-549 and MDA-MB-468 cells were maintained in RPMI (Gibco) supplemented with 10% FBS (Sigma Aldrich), at 37°C in a 5% CO_2_ incubator. MCF-10A were maintained in DMEM-F12 (Gibco) supplemented with 10% FBS (Sigma Aldrich), 5% penicillin-streptomycin (Gibco), 10µg/mL of insulin (Sigma Aldrich), 20ng/mL of EGF (Sigma Aldrich), 100ng/mL of cholera toxin (Sigma Aldrich) and 0.5µg/mL of hydrocortisone (Sigma Aldrich), at 37°C in a 5% CO_2_ incubator.

### DDR1 knock-out (KO) cell line generation

MDA-MB-231 cells KO for DDR1 were generated, using CRISPR-Cas9, by the CRISP’Edit facility in Bordeaux (TBM Core). MDA-MB-231 were transfected by nucleofection with CRISPR/Cas9 protein as well as a crDNA (5’-TGGGAAACACCGACCCTGCGGGG-3’) to deplete DDR1. Transfected cells were then sorted in order to seed 1 cell per well and proceed to clonal selection. Clones were expanded and tested for DDR1 protein expression using western blotting.

### Extracellular matrix (ECM) preparations

For most experiments, cells were seeded on gelatin coverslips coated with a thin layer of collagen. Type I collagen (rat tail, Corning, 354236) and type III collagen (human type III, CellSystems, 5021) were diluted in DPBS containing calcium and magnesium (Dulbecco’s Buffered-Phosphate Saline, Gibco) at a final concentration of 0.5mg/ml. For some immunofluorescence and videomicroscopy experiments, type I collagen was coupled to 5-carboxy-X-rhodamine succinimidyl ester (Fisher Scientific, 11590146). In all cases, collagens were polymerized for 4 h at 37°C before cell seeding. Non-fibrillar collagen matrix was made by diluting type I collagen at 0.5mg/mL in 1M acetic acid (Merck, 100063) and incubated for 4 h at 37°C before cell seeding. Collagen crosslinking was performed using recombinant LOX-L2 (50 ng/mL for 4 days) (2639-AO, R&D system).

Matrigel (Dutscher, 356231) was diluted in PBS (Phosphate Buffer Saline, Gibco) at a final concentration of 0.5mg/mL and polymerized for 2 hours at 37°C before cell seeding. Gelatin-coated coverslips were made using gelatin from Sigma Aldrich (G1393). Coverslips were incubated on 1X gelatin for 20 min before being fixed with 0.5% glutaraldehyde (Electron Microscopy Science) for 40 min at room temperature. Coverslips were washed 3 times with PBS before cell seeding. Fibronectin (F1056, Sigma Aldrich) was used at a final concentration of 1mg/mL, diluted in sterile water, and was polymerized for 45 min at room temperature. Unless stated otherwise, cells were incubated overnight before analysis.

### Inhibition of cell migration

MDA-DDR1-GFP cells were seeded on coverslips coated with type I collagen as previously described. Drugs were added at the time of seeding, as follow: cytochalasin D (C2618, Sigma Aldrich) was added to the cell medium at a concentration of 5µg/mL, blebbistatin (B0560, Sigma Aldrich) was added at a concentration of 50µM. Both drugs were incubated overnight at 37°C. DMSO (D8418, Sigma Aldrich) was used as a control condition.

### Cell detachment

In order to analyze the collagen fibers only, cells were removed from the plate by a wash with warm PBS followed by multiple baths of warm 20mM EDTA (Euromedex, EU0084) for 5 min at 37°C. This allowed to remove the majority of the cells without damaging the collagen and the collagen-tracks. For proteomics and RNA-sequencing analysis, the remaining cells were removed by laser microdissection using PALM type 4 microscope (Zeiss).

### Immunofluorescence and imaging

Cells were fixed using 4% paraformaldehyde (PFA, Electron Microscopy Sciences) for 10 min at room temperature then rinsed 3 times with PBS. Cells were permeabilized with 0.2% Triton X-100 (T9284, Sigma Aldrich) for 10 min at room temperature before being rinsed twice with PBS. Cells were then incubated with primary antibodies (1:100 diluted in 4% BSA-PBS) for 40 min at room temperature, rinsed 3 times with PBS and incubated with secondary antibodies (1:200 diluted in 4% BSA-PBS) for 30 min at room temperature. Nuclei were stained with Hoechst (1:1,000 diluted) and actin was stained with phalloidin (1:200 diluted). For membrane staining, live cells were rinsed once with warm PBS and incubated with Cell Mask (1:2,000 diluted in warm PBS) for 5 min at 37°C before fixation with 4% PFA. Coverslips were mounted on microscope slides using Fluoromount-G mounting media (SouthernBiotech) and were imaged using SP5 confocal microscope (Leica). Collagen is observed by internal reflection microscopy (IRM)^41^. Images were analyzed using ImageJ or Fiji softwares.

### Videomicroscopy

MDA-DDR1-GFP cells were seeded in a bottom-glass dish (µ-Dish 35mm, Ibidi) on top of a layer of fluorescent type I collagen. After 4 h, cells were imaged every 4 min for 15 h using W1 LiveSR spinning disk microscope at the Bordeaux Imaging Center. Videos were then reconstituted using ImageJ. SPI-488, SPI-561 and SPI-DIC were used to image collagen-tracks, collagen and transmission respectively.

### Electron microscopy

#### SEM

MDA-DDR1-GFP cells were seeded on coverslips previously coated with type I collagen. The day after, coverslips were fixed with 4% PFA and an immunofluorescence experiment was performed as described before. Coverslips were imaged using SP5 confocal microscope (Leica) before being processed for SEM. Cells were fixed with 2.5% glutaraldehyde (electron microscopy) and 4% PFA in a phosphate saline buffer for 30 min at room temperature then overnight at 4°C. They were further rinsed twice in 0.1M Phosphate Buffer (Electron Microscopy Sciences, 50-365-831) and dehydrated in a series of increasing ethanol concentrations (30 –100%). Samples were dried via critical point drying (Leica EM CPD300, Austria). Afterwards the cover glass was mounted onto specific stubs and coated with platinum, using a sputter coater (Q150T, Quorum Technologies, Kent, UK). Observations were done at 2 kV, in "high vacuum" mode, with a GeminiSEM 300 FESEM (ZEISS Germany), at the Bordeaux Imaging Center.

#### TEM

Analysis of collagen-tracks by TEM was adapted from Croissant et al^26^. Briefly, MDA-DDR1-GFP cells were cultured in a 35-mm glass bottom dish equipped with a square-patterned coverslip (MatTek, P35G-1.5-14-C-GRID) for 24 h. The day after, coverslips were fixed with 4% PFA and an immunofluorescence was performed as previously described. Coverslips were imaged using SP5 confocal microscope (Leica). They were then post-fixed overnight at 4°C in a mixture of 4% PFA and 2% glutaraldehyde in 0.1 M cacodylate buffer (pH 7.4). They were treated with 1% osmium tetroxide in 0.1 M cacodylate buffer for 10 min at room temperature, dehydrated with ethanol and finally embedded in Epon-Araldite. 70nm sections were collected using the EM UC7 ultramicrotome (Leica, Wetzlar, Germany) and stained successively with 5% uranyl acetate and 1% lead citrate. TEM observation was performed with a FEI CM120 operated at 120 kV. Images were recorded with a USC1000 slow scan CCD camera (Gatan, Pleasanton, CA, USA).

### Second harmonic generation (SHG) microscopy

The confocal microscope was a Leica TCS SP5 on an upright stand DM6000 (Leica Microsystems, Mannheim, Germany), using objectives HCX Plan Apo CS 63X oil NA 1.40. For confocal microscopy the lasers used were Argon 488 nm, 561 nm and 896 nm. The multiphoton microscopy was done with pulsed laser Mai Tai HP (Spectra-Physics, Irvine, USA) tuned at 896 nm. SHG imaging has been done on the transmitted light PMT with 2 filters on the pathway to specifically detect the SHG signal: a 680 nm shortpass filter and a 448/20 bandpass filter.

### Western blotting

Cells were washed once with ice-cold PBS, scraped into ice-cold RIPA lysis buffer (0.1% SDS, 1% NP-40, 150mM NaCl, 1% sodium deoxycholate, 25mM Tris-HCl pH 7.4, Complete and PhoStop inhibitors (Roche)) and lysed for 30 min at 4°C. Lysates were clarified by centrifugation at 13,000g for 15 min at 4°C. Protein concentrations were determined using Lowry reagent (DC^TM^ Protein Assay, BioRad), according to the manufacturer’s protocol. Lysates were boiled at 95°C for 10 min and separated by SDS-PAGE electrophoresis on 10% acrylamide gels (Fast Cast kit, BioRad) at 100V for 75 min. Samples were transferred onto nitrocellulose membranes using a Transblot transfer system (BioRad) and proteins were revealed using Ponceau S solution (Sigma Aldrich). Membranes were blocked using Odyssey Blocking Buffer (Odyssey) for 1 h at room temperature before being incubated with primary antibodies (1:1,000 diluted in 5% BSA-1XTBS-0.2%Tween) (TBS 10X, Euromedex, ET220; Tween, Sigma Aldrich, P7949) overnight at 4°C. The next day, membranes were washed 3 times with TBST and incubated with secondary antibodies (1:5,000 diluted in 5% BSA-TBST) for 1 h at room temperature. Membranes were washed 3 times with TBST before exposure using a Chemidoc system (BioRad).

### RNA extraction and RT-qPCR

MCF-7 cells were incubated for 72 h on collagen, collagen + conditioned medium of MDA-DDR1-GFP or collagen decorated with collagen-tracks from MDA-DDR1-GFP cells. mRNAs were extracted from cultured cells using the Nucleospin RNA kit (Macherey Nagel) according to the manufacturer’s instructions. cDNA was synthesized from 1µg of total RNA with Maxima Reverse Transcriptase (Fermentas). 5ng of cDNA were then subjected to PCR amplification on an RT-qPCR system using the CFX96 Real Time PCR detection system (Biorad). The SYBR® Green SuperMix for iQTM (Quanta Biosciences, Inc.) was used with the following PCR amplification cycles: initial denaturation (95°C for 10 min), followed by 40 cycles of denaturation (95°C for 15 s) and extension (60°C for 1 min). Results are expressed in Fold change according to the 2-ΔΔCT method-. 18S ribosomal RNA was used as internal controls. All primers used are listed in table 1.

### Mass spectrometry

#### Sample preparation (cell preparation)

Two conditions were compared: a decellularized ECM decorated with collagen-tracks to a control ECM incubated with conditioned medium from the same cells. For the track-containing sample, 140,000 MDA-DDR1-GFP cells were seeded on a lumox® 50 mm dish covered with a silicon ring membrane (Sarstedt) previously coated with 0.5mg/ml type I collagen (Corning, prepared as previously described). The control dish was coated with 0.5mg/ml type I collagen incubated with conditioned media from the same MDA-DDR1-GFP cells. After an overnight incubation, cells were detached using 12 to 16 baths of EDTA 20mM for 5 min at 37°C. The remaining cells were removed by laser microdissection using PALM type 4 microscope (Zeiss) in an adhesive cap. The ECM were then lysed into an ice-cold RIPA lysis buffer.

#### Sample preparation (for mass spectrometry)

Three independent biological replicates on protein extracts from collagen-tracks and a control have been performed. Proteins were loaded on a 10% acrylamide SDS-PAGE gel and proteins were visualized by Colloidal Blue staining. Migration was stopped when samples had just entered the resolving gel and the unresolved region of the gel was cut into only one segment. Each SDS-PAGE band was cut into 1 mm x 1 mm gel pieces. Gel pieces were destained in 25mM ammonium bicarbonate (NH_4_HCO_3_), 50% Acetonitrile (ACN) and shrunk in ACN for 10 min. After ACN removal, gel pieces were dried at room temperature. Proteins were first reduced in 10mM dithiothreitol, 100mM NH_4_HCO_3_ for 60 min at 56°C then alkylated in 100mM iodoacetamide, 100mM NH_4_HCO_3_ for 60 min at room temperature and shrunk in ACN for 10 min. After ACN removal, gel pieces were rehydrated with 50mM NH_4_HCO_3_ for 10 min at room temperature. Before protein digestion, gel pieces were shrunk in ACN for 10 min and dried at room temperature. Proteins were digested by incubating each gel slice with 10ng/µl of trypsin (V5111, Promega) in 40mM NH_4_HCO_3_, rehydrated at 4°C for 10 min, and finally incubated overnight at 37°C. The resulting peptides were extracted from the gel by three steps: a first incubation in 40mM NH_4_HCO_3_ for 15 min at room temperature and two incubations in 47.5% ACN, 5% formic acid for 30 min at room temperature.

#### NanoLC-MS/MS analysis

The three collected extractions were pooled with the initial digestion supernatant, dried in a SpeedVac, and resuspended with 0.1% formic acid. NanoLC-MS/MS analysis were performed using an Ultimate 3000 RSLC Nano-UPHLC system (Thermo Scientific, USA) coupled to a nanospray Orbitrap Fusion™ Lumos™ Tribrid™ Mass Spectrometer (Thermo Fisher Scientific, California, USA). Each peptide extracts were loaded on a 300 µm ID x 5 mm PepMap C_18_ precolumn (Thermo Scientific, USA) at a flow rate of 10 µL/min. After a 3 min desalting step, peptides were separated on a 50 cm EasySpray column (75 µm ID, 2 µm C_18_ beads, 100 Å pore size, ES803A rev.2, Thermo Fisher Scientific) with a 4-40% linear gradient of solvent B (0.1% formic acid in 80% ACN) in 48 min. The separation flow rate was set at 300nL/min. The mass spectrometer operated in positive ion mode at a 2.0 kV needle voltage. Data was acquired using Xcalibur 4.1 software in a data-dependent mode. MS scans (m/z 375-1500) were recorded at a resolution of R = 120000 (@ m/z 200) and an AGC target of 4×10^5^ ions collected within 50 ms, followed by a top speed duty cycle of up to 3 s for MS/MS acquisition. Precursor ions (2 to 7 charge states) were isolated in the quadrupole with a mass window of 1.6 Th and fragmented with HCD@30% normalized collision energy. MS/MS data was acquired in the ion trap with rapid scan mode, AGC target of 3×10^3^ ions and a maximum injection time of 35 ms. Selected precursors were excluded for 60 s.

#### Databases and result analysis

Protein identification was done in Proteome Discoverer 1.4. Mascot 2.5 algorithm was used for protein identification in batch mode by searching against a UniProt *Homo sapiens* protein database (74,489 entries, released May 16, 2019 https://www.uniprot.org/ website). Two missed enzyme cleavages were allowed for the trypsin. Mass tolerances in MS and MS/MS were set to 10 ppm and 0.6 Da. Oxidation (M) and acetylation (K) were searched as dynamic modifications and carbamidomethylation (C) as static modification. Raw LC-MS/MS data were imported in Proline Web for feature detection, alignment, and quantification^48^. Protein identification was accepted only with at least 2 specific peptides with a pretty rank=1 and with a protein FDR value less than 1.0% calculated using the “decoy” option in Mascot. Label-free quantification of MS1 level by extracted ion chromatograms (XIC) was carried out with parameters indicated previously^49^. The inference of missing values was applied with 5% of the background noise. A protein enrichment in the collagen-tracks was considered if the ratio of collagen with cell condition vs. collagen without cell condition was greater than or equal to 2. A two-tailed paired t-test was applied to select significant enrichments (p<0.05).

#### Data Availability

The mass spectrometry proteomics data have been deposited to the ProteomeXchange Consortium via the PRIDE^50^ partner repository with the dataset identifier PXD035152.

### RNA and microRNA-sequencing

#### Sample preparation (cell preparation)

For mRNA and microRNA seq of collagen-tracks, the same analytic design was used, we compared a decellularized ECM decorated with collagen-tracks to a control ECM incubated with conditioned media from the same cells. For the track-containing sample, 140,000 MDA-DDR1-GFP cells were seeded on 4 silicone dishes previously coated with 0.5mg/ml type I collagen (Corning, prepared as previously described). The control dish was coated with 0.5mg/ml type I collagen incubated with conditioned media from the same MDA-DDR1-GFP cells. After an overnight incubation, cells were detached using 12 to 16 baths of 20mM EDTA for 5 min at 37°C. The remaining cells were removed by laser microdissection using PALM type 4 microscope (Zeiss). The decellularized ECM were then lysed with 300µl Trizol and kept at -80°C before being used. For MCF-7 cells internalization mRNA-seq, MCF-7 cells were seeded on collagen with collagen-tracks or not for 72h RNA extraction was performed using

#### RNA extraction

RNA was extracted using the Direct-zol RNA microprep kit (Zymo Research) following manufacturer’s instructions, RNA purity was assessed by capillary electrophoresis (LabChip RNA pico sensitivity assay, Perkin Elmer). For MCF-7 cells internalization mRNA-seq RNA extraction was peformed using Nucleospin RNA kit (Macherey Nagel).

#### RNA seq library synthesis

Small-RNA-seq libraries were synthesized using NEXTflex Small-RNA Seq kit v3 (Perkin Elmer) following the manufacturer’s instructions. Briefly, 3’ adenylated adapters, followed by 5’ adapters were sequentially added to RNA strands. Then, RNA samples were reverse transcribed using the 3’ adaptations as the template for the RT primer. Finally, cDNAs were amplified by PCR. Laser microdissection resulted in a low amount of RNA extracted, therefore RNA-seq libraries were synthesized with an adapted Smart-seq protocol adapted to low input^51^. Briefly, mRNA were reverse transcribed with an oligo-dT primer containing an adapter sequence and double-stranded cDNA was obtained with a template switching oligo (TSO) containing the same adapter sequence as the oligo-dT. After a pre-amplification with 15 cycles of PCR, we used the NEXTflex Rapid XP DNA-seq kit v2 (Perkin Elmer) to synthesize RNA-seq libraries. Before sequencing, libraries were qualified by capillary electrophoresis (LabChip GX, Perkin Elmer) and quantified by qPCR (NEBNext Library Quant Kit for Illumina, New England Biolabs). Sequencing was performed on a NextSeq 2000 system (Illumina) in a 2×100bp mode. Oligo-dT sequence: 5’-AAGCAGTGGTATCAACGCAGAGTAC(T)_30_VN. TSO sequence: 5′-AAGCAGTGGTATCAACGCAGAGTACrGrG+G, in which rG indicates riboguanosines and +G indicates a locked nucleic acid (LNA)-modified guanosine.

#### Identification and quantification of microRNAs and mRNAs by bioinformatic analysis

Concerning collagen-tracks Small-RNA-seq, to obtain clean reads, the adaptor sequences (from the libraries preparation), and the 4 nucleotides inserted between the adaptor sequence and the sequence of the microRNAs were removed, using Cutadapt tool (version 1.18^52^). The Human genome reference was downloaded from the UCSC Genome Browser (GRCh38 Assembly). microRNA identification and quantification were performed using the program miRPro^53^, which uses the main algorithm of the software miRDeep2^54^. The main advantage of miRPro compared to miRDeep2 is that it proposes consistent and unified names for novel precursors and their mature miRNAs in all libraries of the samples. The reads issued from the sequencing are aligned against the genome of reference by using Novoalign^55^. Sequences matching known Human small RNAs such as rRNA, scRNA, snoRNA, snRNA and tRNA or degradation fragments of mRNAs were excluded in further analysis, and only sequences that perfectly matched the Human genome along their entire length and recognized as miRNAs were quantified.

Concerning MCF-7 mRNA-seq, adapter sequences were removed using Cutadapt tool (version 1.18^52^), then sequences were further cleaned using the fastp software (default parameters^56^). Clean reads were then aligned to the Human reference genome (GRCh38 Assembly) using the STAR software (v2.7.1a, default parameters^57^) and quantified using the quant-mode of the STAR software. Raw counts were used as input in DESeq2 to identify differentially expressed genes between MCF-7 cells grown in presence or not of collagen-tracks. Results were filtered based on p-value < 0.05 and a Log2FC> |1.5| for further analysis such as Gene Ontology and STRING interactome pathway analyses.

#### Integrative analysis

Analysis of biological pathways associated with mRNAs or proteins significantly enriched in collagen-tracks was performed by Gene Set Enrichment Analysis (GSEA) against the GO Molecular Function database. The pathways potentially targeted by the following microRNAs: hsa-let-7f-5p, hsa-miR-98-5p, hsa-miR-23a-3p, hsa-let-7e-5p, hsa-miR-486-5p, hsa-miR-146a-5p, hsa-miR450a-5p were analyzed in the same way by GSEA, having previously identified its potential targets from the mirDB database.

### miRNA inhibitors

miRCURY LNA miRNA Power Inhibitors (1 nmol) (hsa-let-7e-5p ID-YI04101354-DCA, hsa-miR-23a-5p ID-YI04101840-DCA, hsa-miR-98-5p YI04100423-DCA, hsa-miR-486-5p ID-YI04100385-DCA, hsa-miR-146a-5p ID-YI04100676-DCA, hsa-miR-450a-5p ID-YI04101851-DCA, hsa-let-7f-5p ID-YI04100302-DCA) and control (ID-YI00199006) were purchased from Qiagen. 200,000 MCF-7 cells were seeded in 6 well plate in 1.5ml of medium. A mix with miRNA inhibitor or control (5 nM final concentration), 5µl of lipofectamine RNAimax (Invitrogen) and 500µl of serum-free opti-MEM (Fisher Scientific) were prepared, incubated for 15 min at room temperature and added dropwise. After 4 days, cells were seeded on fluorescent gelatin for degradation assay and RNA extraction as previously described.

### 3D collagen invasion assay

500µl of solution was prepared for 3 Boyden chambers following the protocol described by Di Martino *et al.*, 2017^58^. The type I collagen solution was prepared to a final concentration of 2mg/mL. 5-carboxy-X-rhodamine succinimidyl ester was added at a final concentration of 1µg/mL. The mix was incubated for 5 min on ice under the sterile conditions, protected from light. Then, 50µL of PBS 10X and 6,25µL of 1M sodium hydroxide (NaOH) were added to the mix. Sterile water was added to a final volume of 500µL. 100µL of the mix was added to the Boyden chamber and then incubated during 1h at 37°C. 30,000 MDA-DDR1-GFP cells were added on the top of the gel in complete media (DMEM supplemented with 10% FBS). DMEM + 20% FBS were added at the bottom of the gel to be used as a chemoattractant. After 72h, cells were fixed and immunofluorescence was performed as described in the protocol. The gels were imaged by second harmonic generation microscopy.

### Transwell migration assay

Transwell chambers (8.0µm pore size, Falcon 353097) were coated with 150µl of Matrigel (80µg/ml) in a 24-well plate. For each condition, 40,000 MCF-7 cells per transwell were seeded in DMEM deprived of FBS. As a chemoattractant, DMEM containing 20% FBS was used in the lower part of the transwell in the 24-well plate. After 24 h, the media was carefully removed, cells were fixed with 4% PFA for 10 min and stained with crystal violet (Sigma-Aldrich) for 20 minutes at room temperature. Cells in the upper part of the transwell membrane were removed by wiping with a cotton swab. A phase-contrast microscope (Zeiss) was used to image migrated cells in the lower chambers, 3 images per condition and per replicate were acquired. Images were analyzed using ImageJ software.

### ECM degradation assay

Sterile coverslips were coated with gelatin 488 (Oregon Green™ 488 Conjugate Thermofisher G13186) for 20 min, then fixed with 0.5% glutaraldehyde (15960, Electron Microscopy Sciences) for 40 min. MCF-7 cells were seeded on gelatin 488 coated coverslips and incubated for 35 h, before being fixed with 4% PFA for 10 min. Additionally, MDA-DDR1-GFP and MCF-7 cells with collagen-tracks were seeded on gelatin 488 coated coverslips and treated with 25µM GM6001 (MMP inhibitor) or with DMSO as control. Cells were stained with hoechst as previously described. Coverslips were imaged under the epifluorescence microscope (Zeiss) using the 40X oil immersion objective. A total of 10 images were acquired for each condition, and experiments were done in three biological replicates. The area of degradation was quantified using ImageJ and normalized to the number of nuclei in each image.

### Adhesion activation assay

MDA-MB-231 and MDA-DDR1-GFP cells were seeded on coverslip coated with type I collagen as described above. 2 h after seeding, cells were treated with 0.5mM of manganese (Mn2+) or with water (control). Cells were fixed 6 h after treatment and the immunofluorescence was performed as previously described. A total of 5 images were performed for each condition using confocal microscope (SP5 Leica) and collagen-tracks were quantified using ImageJ and normalized to the number of nuclei.

### In vivo xenografts

#### Cell preparation

MDA-DDR1-GFP cells were transduced (multiplicity of infection 10) with lentiviral particles generated by the Vect’Ub facility (TBM Core, Bordeaux) from the pHR-PP7-mCherry-Caax lentiviral plasmid (#74925, Addgene). The final cells were named MDA-DDR1-GFP-Caax-mCherry.

#### Xenografts

MDA-DDR1-GFP-Caax-mCherry cells were trypsinized and resuspended at 4 million cells in 100µl media without FBS. 4 million cells were intradermally injected into left and right flanks of 3 NSG (NOD/LtSz-scid IL2Rγnull) anesthetized mice. Mice were sacrificed 15 days after injection in order to get primary tumor development as well as local invasion. Primary tumors and surrounding tissues were collected and snap-frozen in OCT (Tissue-Tek). Six 7µm and one 50µm serial slices were done using the Cryostat (Leica). 7 µm slices were mounted on microscope slides while 50µm slices were fixed using 4% PFA for 10 min and kept in 1X PBS before use. All animal experiments were conducted in accordance with protocols approved by the French Experimental Animal Care Commission.

### Immunohistochemistry

Immunohistochemistry was performed with the help of the histopathological facility in Bordeaux (TBM Core).

#### 7µm slices

Half of the 7µm slices were stained with anti-GFP and anti-mCherry antibodies while the other half were stained with hematoxylin, eosin and saffron (HES). For the GFP-mCherry staining, slides were permeabilized for 10 min in 1X TBST before automated staining with Dako-Agilent autostainer. Slides were first incubated with anti-GFP (ab6673, Abcam) and anti-mCherry (ab183628, Abcam) primary antibodies for 1 h at room temperature. After two washes, slides were incubated with secondary antibodies (anti-rabbit 594 #DI-1794, Vectafluor and anti-goat 488 #DI-3788, Vectafluor) and Hoechst for 40 min at room temperature. After washes, the slides were mounted with Fluoromont-G mounting media (SouthernBiotech).

HES staining was performed using a Leica ST stainer. Slides were incubated in two baths of toluene for 5 min each, then dehydrated into two successive baths of 100% ethanol for 1 min, then into 95% ethanol for 1 min. After 3 min of hydration in water, slides were stained with Mayer hematoxylin for 3 min, then rinsed with water for 3 min. Slides were then stained with 1% aqueous eosin then rinsed with water for 3 min. After 2 successive baths of 100% ethanol for 15 s, slides were stained with 2% safran before being dehydrated by 2 baths of 100% ethanol. Slides were then incubated into 2 successive baths of toluene for 5 min before being mounted using Eukitt mounting media (Sigma). All of the 7µm slices were imaged using a Hamamatsu Nanozoomer 2.0HT at the Bordeaux Imaging Center, University of Bordeaux.

### 50µm slices

After fixation, the desired 50µm slices were stained with anti-GFP and anti-mCherry antibodies. Briefly, slides were incubated overnight in TBST for permeabilization. They were then incubated with primary antibodies for 48 h (1:200 diluted in TBST), rinsed 3 times with PBS (1 day wash) and incubated for 24 h with secondary antibodies (1:500 diluted in TBST). After 3 more washes, slices were mounted on microscope slides using the Fluoromont-G mounting media. Slides were imaged using a Leica DM6000 TCS SP5 MP microscope (Bordeaux Imaging Center, Bordeaux).

### Primary cell collection and immunofluorescence

Triple negative breast cancer primary tumor from a surgical resection was digested for 3 h with collagenase then injected into the mammary ducts of a female NSG mouse (Jackson Strain 005557; passage 1). After 9 weeks, the resulting PDX was harvested, dissociated with collagenase and the cells were again injected into the mammary ducts of NSG mice (passage 2). After 16 weeks, the resulting PDX was harvested, digested with collagenase and frozen in cryo3 (StemAlpha, Lyon). After thawing, the tissue organoids were broken up by vigorous pipetting and debris was removed by extensive washing. The remaining material was in clumps of 10-20 cells that were infected overnight with 250µl LKB virus (vDRM166, luciferase-mKate2-blasticidin, Addgene #183502; titre 1×10^8^/ml) in a final volume of 500µl organoid medium^38^. The cells were then rinsed with organoid medium and injected into the mammary ducts of 16-week-old female NSG mice (passage 3). Tumor growth was followed by bioluminescence and the mouse was sacrificed 12 weeks after injection. At sacrifice the gland was thickened with tumor infiltrating the entire gland and mechanically dissociated using a razor blade. Dissociated samples were digested during one hour in DMEM with Collagenase Type I, 1mg/mL (Sigma-Aldrich). Samples were cleaned on 100μm and 40μm filters. Cells were cultivated in Advanced DMEM (Gibco), 5% FBS, HEPES (Gibco), Glutamax (Gibco), 10ng/mL human EGF (Peprotech), 1µg/mL hydrocortisone (Sigma-Aldrich), 10µM Y-27632 (Sigma-Aldrich) and 5µg/mL blasticidin (Sigma-Aldrich)^38,39^. Cells were then seeded on a collagen matrix and immunofluorescence was performed as described above.

### Statistical analysis

Data are reported as the mean ± SEM of at least three experiments. Statistical significance (p<0.05 or less) was determined using analysis of variance (ANOVA) (one way followed by a bonferroni correction), unpaired or paired student t-test as appropriate and performed with GraphPad Prism software. Significance levels are shown as stars.

## Acknowledgments

We are grateful to different facilities core, laser-microdissection (Neurocentre Magendie), Bordeaux Imaging Center (BIC) (Univ. Bordeaux, CNRS *UMS* 3420, INSERM US04) and platforms of the unit TBMCore (US 005), Oncoprot, Histopathology, Vect’UB, CrispEdit. We also thank Pr M. Teichman who provided several breast cancer cell lines.

## Author contributions

Conceptualization: FS, MR

Methodology: LN, LR, MR, ER, NA, SDT, CD AAR, JWD, GMG, NDS, AB, ST, AF

Investigation: LN, LR, MR, ER, NA, RI

Visualization: LN, LR, MR

Supervision: FS

Writing—original draft: LN, LR, MR, FS, VM

## Data and materials availability

All data supporting the findings of this study are available from the corresponding author upon reasonable request. The mass spectrometry data are available via ProteomeXchange with the identifier PXD035152. Source data are provided with this paper.

---

**Figure. S1:**
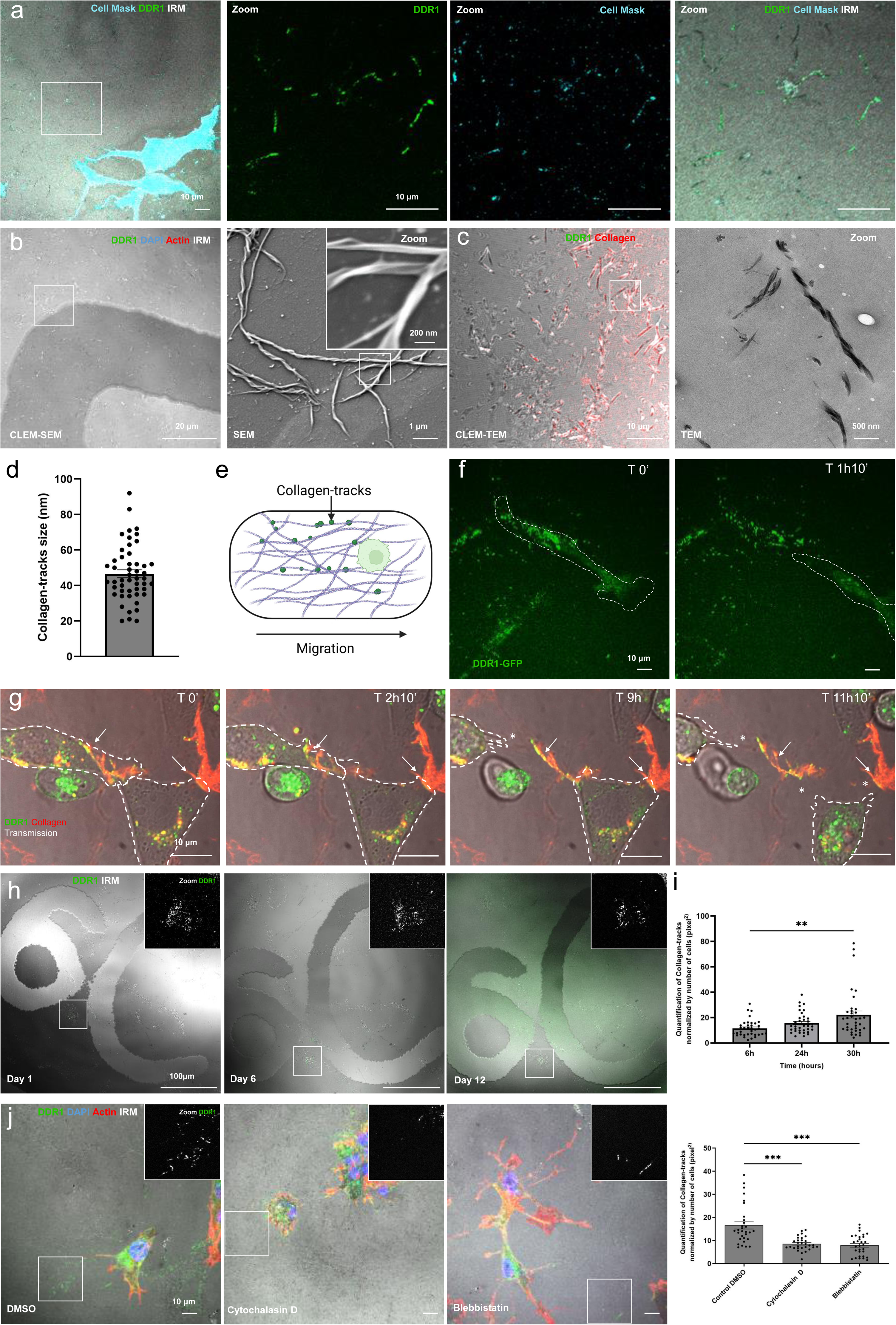
Membrane vesicles containing DDR1 are deposited on collagen fibers during breast cancer cell migration in 2D and 3D. **(a)** Immunofluorescence images of MDA-DDR1-GFP cells seeded on type I collagen and imaged for cell membrane (Cell Mask, in cyan) and DDR1 (in green). Scale bars, 10μm. **(b)** CLEM-SEM images of conditioned media incubated on type I collagen. DDR1 is stained in green, actin in red, nuclei in blue and collagen is imaged by IRM. Scale bars, 20 μm, 1 μm and zoom 200 nm. **(c)** CLEM-TEM images of conditioned media on type I collagen. DDR1 is stained in green and collagen in red for immunofluorescence images. Scale bars: 10 µm and 500 nm. **(d)** Quantification of collagen-tracks size based on SEM microscopy images. Values represent the mean +/- SEM of n=1 experiment with 50 images. **(e)** schematic representation of collagen-track formation during cell migration. **(f)** Images extracted from time lapse imaging (Movie 1) showing collagen-tracks formation after MDA-DDR1-GFP cells migration along type I collagen. Cells were imaged by videomicroscopy for 15h. DDR1 is stained in green. Scale bar, 10 µm. The dash line surrounds the cell to show cell migration. **(g)** Images extracted from time lapse imaging (Movie 2) showing collagen-tracks formation by MDA-DDR1-GFP cells along type I collagen. Cells were imaged by videomicroscopy for 15h. DDR1 is stained in green and collagen in red. The dash lines surround the two cells, white arrows show collagen-tracks on collagen and white stars show the absence of contact between cells and collagen fibers. Scale bar, 10 µm. **(h)** Lifespan of collagen-tracks on collagen fibers without cells. Collagen-tracks were imaged by confocal microscopy on gridded coverslip at day 1, 6 and 12. DDR1 is stained in green and collagen is imaged by IRM. Scale bar, 100 μm. **(i)** Kinetic of collagen-tracks formation over the time by MDA-DDR1-GFP cells on type I collagen. Values represent the mean +/- SEM of n=3 independent experiments (12 images per condition and per replicate) and were analyzed using One-way ANOVA test followed by bonferroni test p<0.001. **(j)** Images and associated quantification of collagen-tracks formation by MDA-DDR1-GFP cells on type I collagen and treated with inhibitors of migration. Cells were imaged by confocal microscopy. DDR1 is stained in green, actin in red, nuclei in blue and collagen is imaged by IRM. Scale bar, 10 µm. Values represent the mean +/- SEM of n=3 independent experiments (10 images per condition and per replicate) and were analyzed using One-way ANOVA test followed by bonferroni test p<0.001.

**Figure S2:**
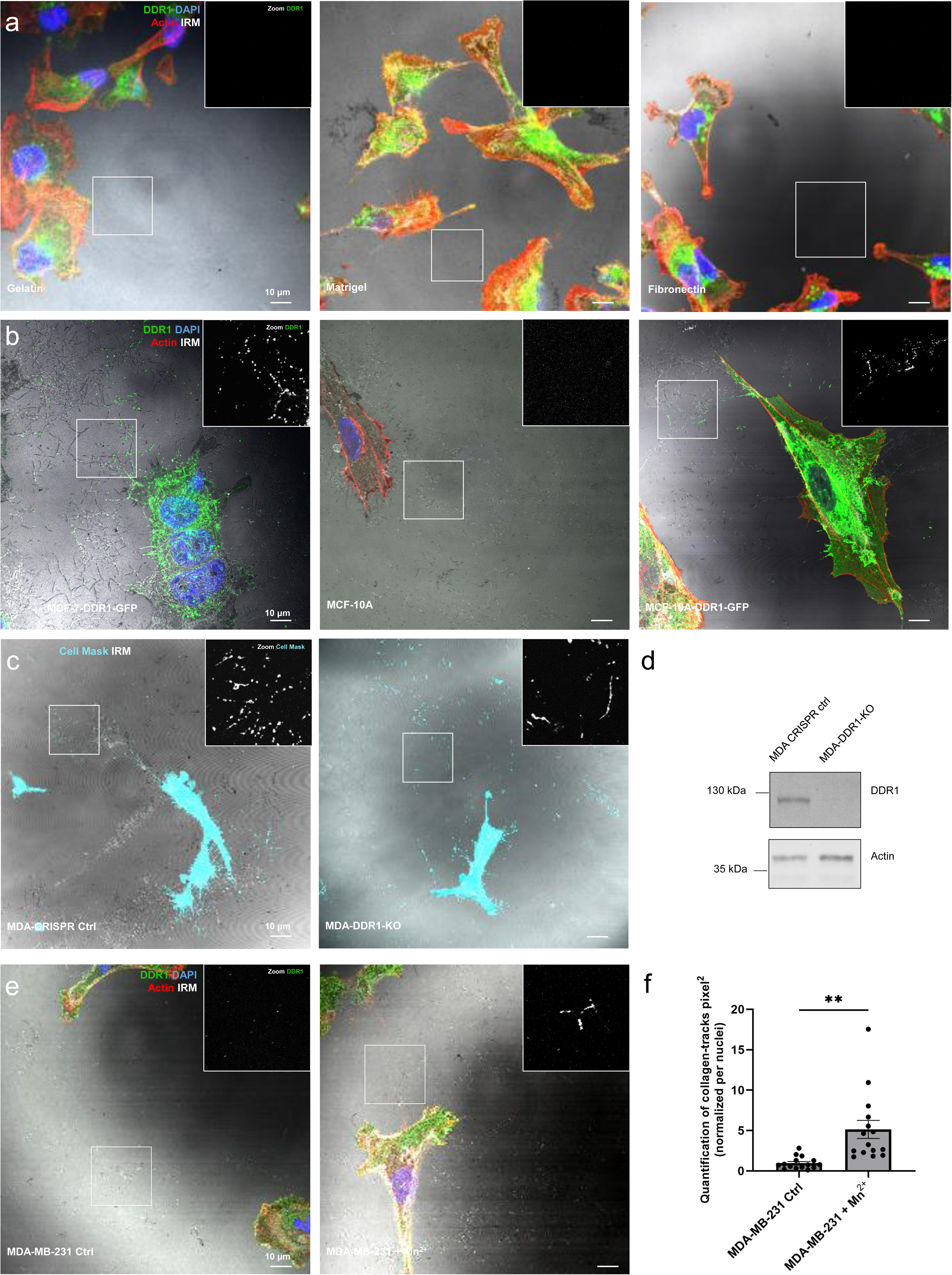
Collagen-track formation is stimulated by Collagen fibers-DDR1/integrin interaction. **(a)** Formation of collagen-tracks on gelatin, matrigel or fibronectin matrices by MDA-DDR1-GFP cells. Cells were imaged by confocal microscopy. DDR1 is stained in green, actin in red, nuclei in blue and collagen is imaged by IRM. Scale bar, 10 µm. **(b)** Immunofluorescence images of collagen-tracks made by MCF-7 overexpressing DDR1 and MCF-10A (WT or overexpressing DDR1) seeded on type I collagen. DDR1 is stained in green, actin in red, nuclei in blue and collagen is imaged by reflection. Scale bar, 10 µm. **(c)** Immunofluorescence images of MDA-MB-231 CRISPR control cells and MDA-MB-231 knockout (KO) cells for DDR1. Collagen-tracks are stained with Cell Mask in cyan and collagen is imaged by IRM. Cells were imaged by confocal microscopy. Scale bar, 10 µm. **(d)** Western Blot analysis of endogenous DDR1 knockout expression in MDA-MB-231 cells. GAPDH was used as the loading control. Experiments were performed in triplicates. **(e)** Immunofluorescence images of collagen-track formation by MDA-MB-231 cells on type I collagen and treated or not with Mn2+. DDR1 is stained in green, actin in red, nucleus in blue and collagen imaged by IRM in grey. Scale bar, 10 µm. **(f)** Quantification of collagen-track formation by MDA-MB-231 cells on type I collagen and treated or not with Mn2+. Values represent the mean +/- SEM of n=3 experiments (5 images per experiment and per condition).

**Fig. S3:**
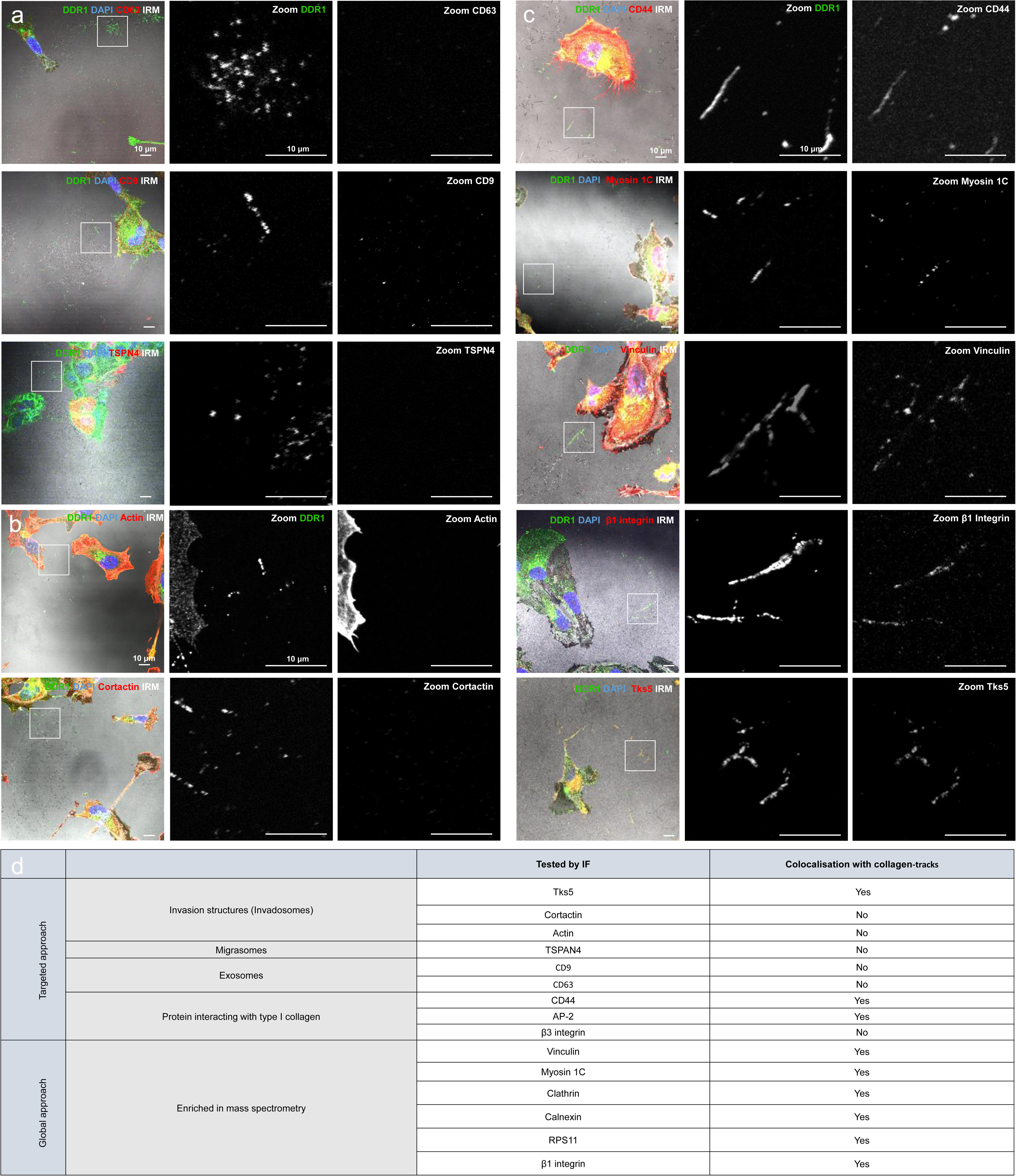
Protein composition reveals collagen-tracks as novel structures. **(a)** Immunofluorescence images of MDA-DDR1-GFP cells seeded on type I collagen and stained for CD63, CD9 or TSPN4 (in red) and DDR1 (in green). Scale bar, 10µm. **(b)** Immunofluorescence images of MDA-DDR1-GFP cells seeded on type I collagen and stained for actin or cortactin (in red) and DDR1 (in green). Scale bar, 10µm. **(c)** Immunofluorescence images of MDA-DDR1-GFP cells seeded on type I collagen and stained for CD44, Myosin 1C, Vinculin, β1 Integrin or Tks5 (in red) and DDR1 (in green). Scale bar, 10 µm. **(d)** Summary table for tested proteins of targeted and global approaches by immunofluorescence.

**Fig. S4:**
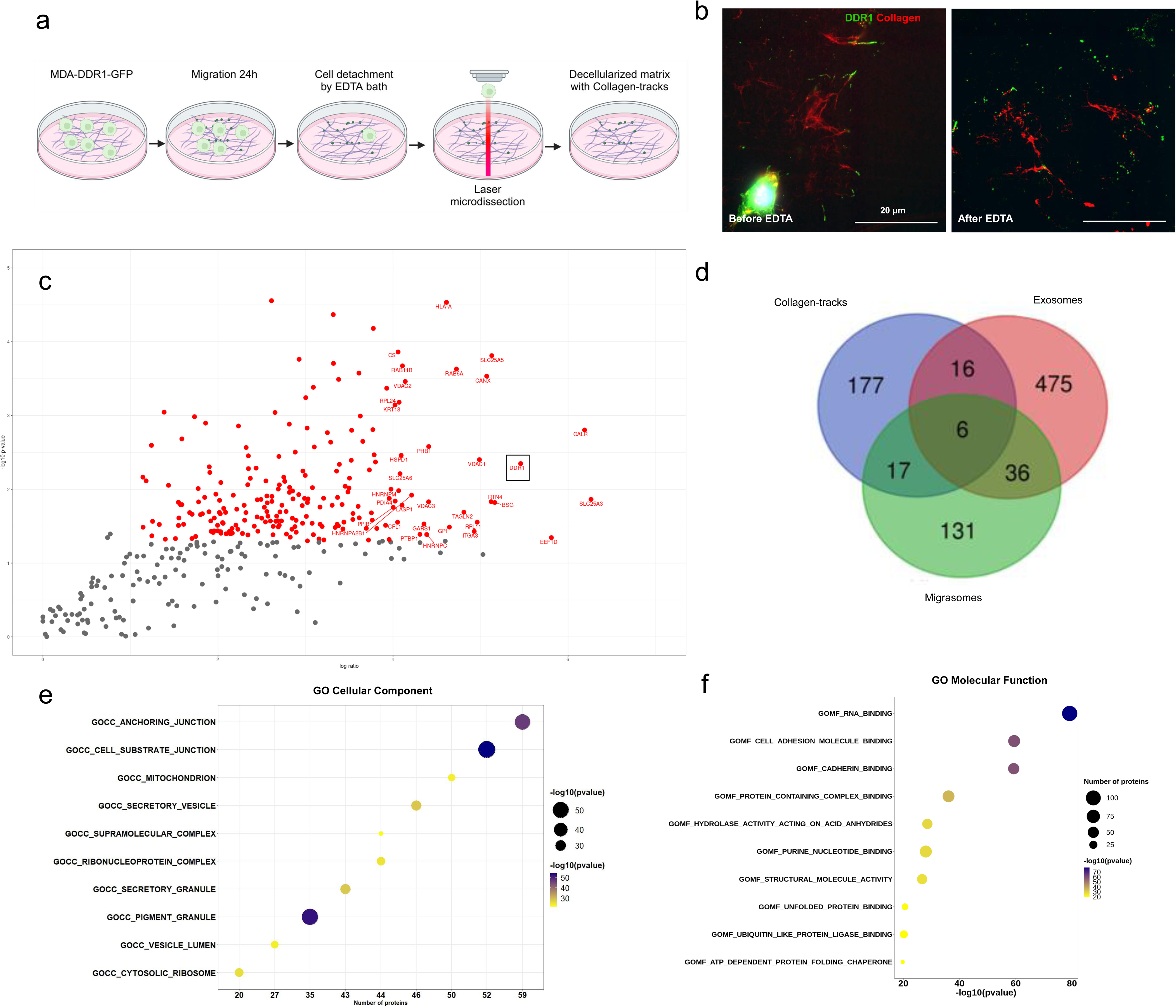
Protein composition reveals collagen-tracks as novel structures. **(a**) Experimental workflow of collagen-tracks formation and decellularization process before mass spectrometry analysis. **(b)** Immunofluorescence images of cell detachment combining EDTA baths and laser microdissection to obtain a fully decellularized matrix decorated with collagen-tracks. DDR1 is stained in green and collagen in red. Scale bar, 20 µm. **(c)** Volcano map highlighting the most expressed proteins enriched in collagen-tracks after mass spectrometry analysis on collagen-tracks. **(d)** Venn diagram comparing collagen-tracks proteome to exosomes and migrasomes proteome extracted from IPA data bank. **(e)** Bubble plot of cellular component of Gene Ontology (GO) enrichment analysis. **(f)** Bubble plot of molecular function of Gene Ontology (GO) enrichment analysis.

**Figure S5:**
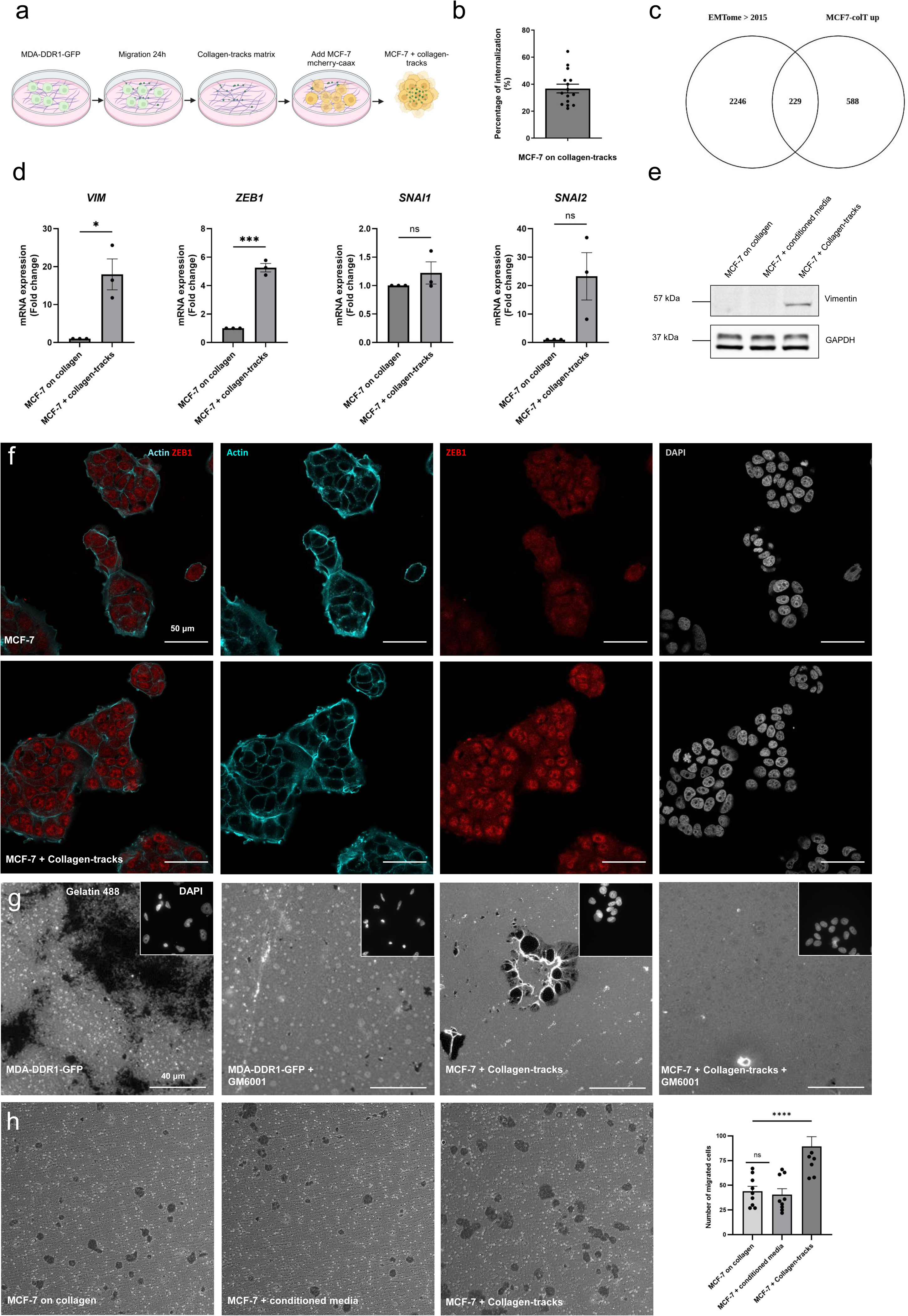
Internalization of collagen-tracks induces a partial-EMT and an invasive switch in recipient cells. **(a)** Experimental workflow of MCF-7 cells internalizing collagen-tracks. **(b)** Quantification of the percentage of MCF-7 cells with internalized collagen-tracks. Values represent the mean +/- SEM of n=3 independent experiments (5 images per replicate). **(c)** Venn diagram comparing MCF-7 collagen-tracks EMT transcriptomic signature to EMTome transcriptomic signature. **(d)** mRNA expression levels of EMT genes in MCF-7 cells on collagen-tracks or collagen only. The graph shows the quantification of *VIM, ZEB1, SNAI1* and *SNAI2* mRNA expression levels normalized to 18S (Fold change). Values are expressed as the mean ± SEM of n=3 independent experiments. Statistics compare all conditions to the collagen condition. A t-test was performed, *p=0.0140, ***p=0.0001. **(e)** Western blot analysis of Vimentin expression in MCF-7 cells on collagen, conditioned media or collagen-tracks. GAPDH was used as the loading control. **(f)** Immunofluorescence images of MCF-7 cells seeded on collagen or collagen-tracks and stained for Zeb1 (in red). Cells were imaged by confocal microscopy. Scale bar, 40 μm. **(g)** Effect of the metalloprotease inhibitor GM6001 on the degradation properties of MCF-7 cells that have internalized collagen-tracks. Representative fluorescent images of microscopic fields where MCF-7 + collagen-tracks cells are treated with or without 20 μM GM6001 inhibitor, MDA-DDR1-GFP are used as positive control. Scale bar 40 μm. **(h)** Quantification of invasion properties of MCF-7 cells into a transwell after incubation on collagen, conditioned media or collagen-tracks with corresponding images can be found on the right. Values represent the mean +/- SEM of n=3 independent experiments and were analyzed using One-way ANOVA test followed by bonferroni test p<0.0001. A total of 3 images were performed for each condition using a confocal microscope.

**Figure S6:**
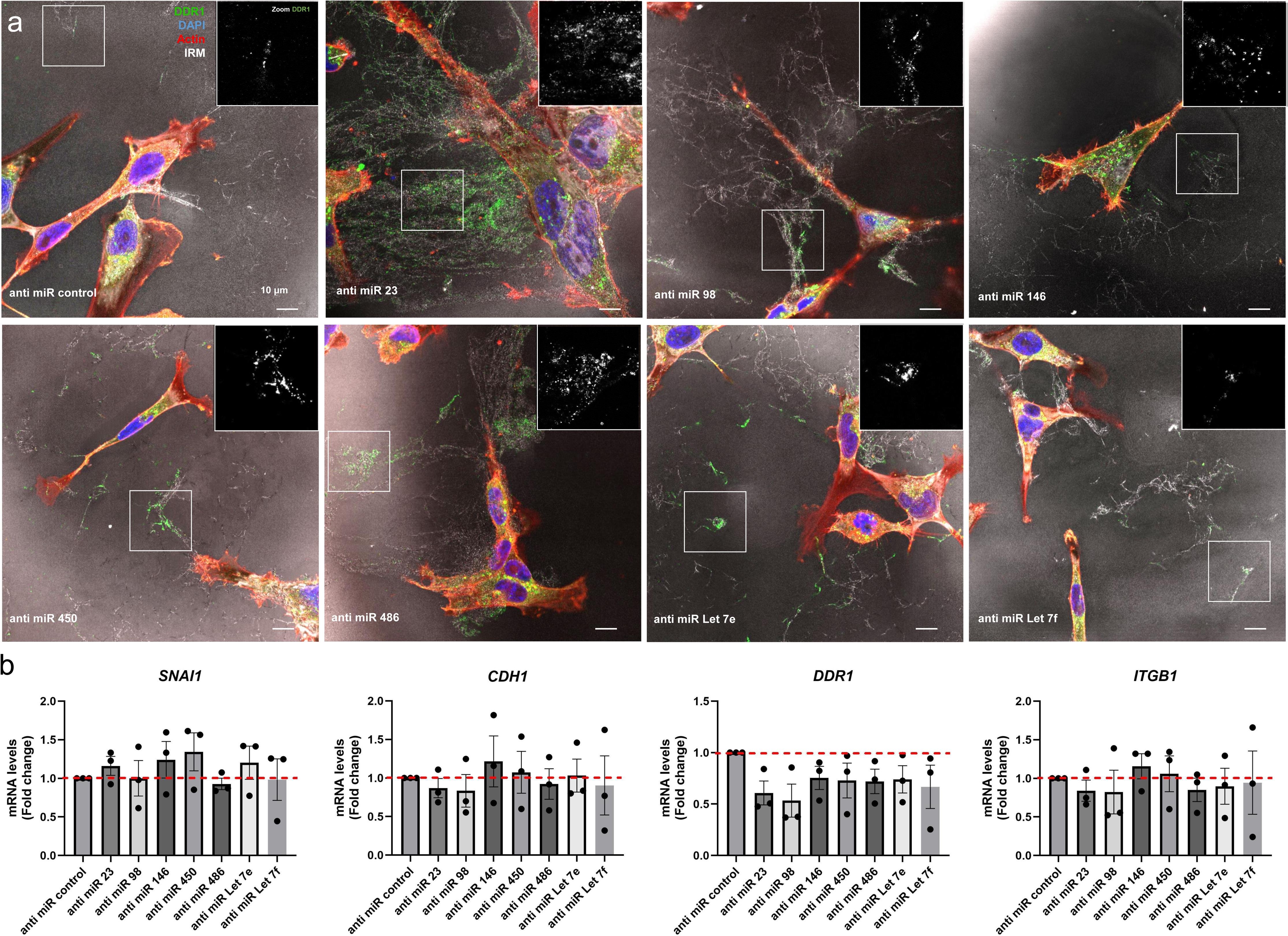
microRNAs contained in collagen-tracks promotes EMT of recipient cells. **(a)** Immunofluorescence images of collagen-tracks after anti-miRNAs or anti-miRNA control treatment in MDA-DDR1-GFP cells. Cells were imaged by confocal microscopy. DDR1 is stained in green, actin in red, nuclei in blue and collagen is imaged by IRM in grey. Scale bar, 10 μm**. (b)** mRNA expression levels of EMT genes in MCF-7 cells after internalization of collagen-tracks previously formed with anti-miRNAs or anti-miRNA control. The graph shows the quantification of *CDH1*, *DDR1,* S*NAIL and ITGB1* mRNA expression levels normalized to 18S (Fold change). Values are expressed as the mean ± SEM of n=3 independent experiments and were analyzed using One-way ANOVA test p<0.001.

**Table S1: Proteomique signature of collagen-tracks**

**Table S2: Summary table of proteins represented in the Venn diagram comparing Collagen-tracks, Exosomes and Migrasomes.**

**Table S3: Transcriptomique EMT signature in MCF-7 with collagen-tracks Movie S1. MDA-DDR1-GFP cells deposit collagen-tracks after cell migration.**

MDA-DDR1-GFP cells were allowed to migrate along type I collagen and formed tracks after cell migration. Cells were imaged by spinning disk microscopy every 4min for 15 hours. DDR1 is stained in green.

**Movie S2. MDA-DDR1-GFP cells deposit collagen-tracks on type I collagen.**

MDA-DDR1-GFP cells were allowed to migrate along type I collagen and imaged by spinning disk microscopy every 4min for 15 hours. DDR1 is stained in green, collagen in red and cells are imaged by transmission.

**Movie S3. MCF-7 cells internalize collagen-tracks deposit by MDA-DDR1-GFP cells along type I collagen.**

MCF-7 stably expressed CAAX-mCherry and were allowed to migrate along type I collagen to internalize collagen-tracks of MDA-DDR1-GFP cells. Cells were imaged by spinning disk microscopy every 4min for 15 hours. DDR1 is stained in green.

